# Polygenic variation in sexual investment across an ephemerality gradient in *Daphnia pulex*

**DOI:** 10.1101/2021.06.23.449662

**Authors:** Karen Barnard-Kubow, Dörthe Becker, Connor S. Murray, Robert Porter, Grace Gutierrez, Priscilla Erickson, Joaquin C. B. Nunez, Erin Voss, Kushal Suryamohan, Aakrosh Ratan, Andrew Beckerman, Alan O. Bergland

## Abstract

Species across the tree of life can switch between asexual and sexual reproduction. In facultatively sexual species, the ability to switch between reproductive modes is often environmentally dependent and subject to local adaptation. However, the ecological and evolutionary factors that influence the maintenance and turnover of polymorphism associated with facultative sex remain unclear. To address this basic question, we studied the ecological and evolutionary dynamics of polymorphism in reproductive strategy in a metapopulation of the model facultative sexual, *Daphnia pulex*, located in the southern United Kingdom. We found that patterns of clonal diversity, but not genetic diversity varied with ephemerality. Reconstruction of a multi-year pedigree demonstrated the co-existence of clones that were found to differ in their investment into male production. Mapping of quantitative variation in male production using lab-generated and field-collected individuals identified multiple putative QTL underlying this trait, and we identified a plausible candidate gene. The evolutionary history of these QTL suggests that they are relatively young, and male limitation in this system is a rapidly evolving trait. Our work highlights the dynamic nature of the genetic structure and composition of facultative sex across space and time and suggests that quantitative genetic variation in reproductive strategy can undergo rapid evolutionary turnover.

## Introduction

Reproductive strategy is a central determinant of evolutionary fitness that varies across the tree of life ^1^. Facultative sex is one type of reproductive strategy and is characterized by organisms that switch between asexual and sexual reproduction. In facultatively sexual organisms, this switch is often linked to periods of dormancy ^2^ and dispersal ^3,4^. Switching between asexual and sexual reproduction can have an impact on the evolutionary dynamics of a species, is subject to local adaptation ^5,6^, and is likely to leave characteristic signatures in the genome ^1,7–10^. Despite the prevalence of facultative sex, open questions remain about its reprocussions on the population genetic structure of populations, the influences of ecological factors on the temporal and spatial distribution of reproductive polymorphism, and the origin of genetic polymorphism in reproductive strategy.

The evolutionary dynamics and genetic consequences of facultative sex are influenced by seasonal fluctuations in habitat suitability. Facultatively sexual species generally reproduce asexually during the favorable season and switch to sexual reproduction when conditions deteriorate in order to produce resting embryos that are able to survive adverse conditions ^11–14^. During periods of asexual reproduction, intraspecific competition leads to clonal selection and the erosion of clonal diversity, a phenomenon well documented in natural populations ^15–21^. Thus, clonal diversity and the number of clones participating in sexual reproduction are expected to be negatively related to the duration of the immediately preceding favorable season ^16,18^, with a greater potential for selfing and inbreeding as the duration of the growing season increases. While a single episode of sexual reproduction can restore Hardy Weinburg equilibrium at individual loci after a loss of clonal diversity ^8^, the long-term effects of multiple rounds of potential inbreeding and selfing on genome-wide patterns of genetic variation remain unclear.

Seasonal fluctuations in habitat suitability are also expected to impact the evolution of reproductive strategy. During periods of favorable conditions, the short-term advantage of asexual reproduction should lead to the establishment and rise in frequency of alleles which divert reproductive effort to the production of clonal females ^22–24^, particularly when the production of males comes at a fitness cost ^25,26^. In support of this advantage, non-male production has evolved multiple times in facultative parthenogens ^6,27,28^ and clones with a low investment in sexual reproduction have been observed to increase in frequency across the asexual growing season ^29,30^. However, when the asexual growing season is short or unpredictable, clones with low sexual investment will not have sufficient time to reap the short term benefits of asexual reproduction and clones with high sexual investment will be favored ^5,30–33^. In highly unpredictable environments sexual investment may even become constant and unlinked from environmental stimuli ^2,34^. Thus reproductive strategy is expected to vary among populations of facultative parthenogens as a result of ecology. However, the evolutionary and ecological dynamics influencing the maintenance of polymorphism in reproductive strategy within populations remain elusive.

*Daphnia* are excellent models to address the basic evolutionary and ecological aspects of facultative sex. *Daphnia* are planktonic crustaceans that frequently live in small water bodies prone to drying. *Daphnia* can only survive drying via sexually produced resting eggs. Thus, the duration of asexual reproduction and frequency of enforced sexual reproduction are directly related to ecology. Several species of *Daphnia* exhibit genetic variation in reproductive strategies such as the propensity for females to reproduce sexually and the rate of male production ^6,30,35–41^. Male production has been lost multiple times ^27,28^, and investment into male production has been linked to habitat ephemerality on a continent-wide scale ^6^, but not at the level of single metapopulations ^36^. Notably, in both *D. pulex* and *D. magna*, dominant non-male producing nuclear encoded alleles have been identified at major effect loci ^27,28^ and there is evidence that other loci may also contribute to variation in male production ^6^. Overall, the large range of genetic polymorphism observed for male production, combined with the variation in associated loci across multiple systems, is consistent with a model that male production is a dynamically evolving trait within species subject to constant allelic turnover.

To advance our understanding of the evolutionary dynamics and genomic consequences of facultative sex in relation to ephemerality, we characterized the genetic basis of variation in male production of *D. pulex* across a series of small ponds in the southern region of the United Kingdom (Figure 1A-C). We focused our attention on three “focal ponds” located in Dorset (Figure 1A), each showing different levels of ephemerality. Using whole genome sequencing of hundreds of clones sampled across multiple years as well as lab phenotyping experiments, we asked: 1) How does ephemerality affect the clonal diversity and genetic structure of populations? 2) Is there genetic variation for male production and what is its genetic architecture? and, 3) What is the evolutionary history of these polymorphisms and how are they distributed among populations? We find that the degree of ephemerality has an impact on the genetic structure of the populations, that there is genetic variation in male production that is unique from previously identified systems, and that this variation is relatively young, suggesting rapid genetic turnover.

**Figure 1.**
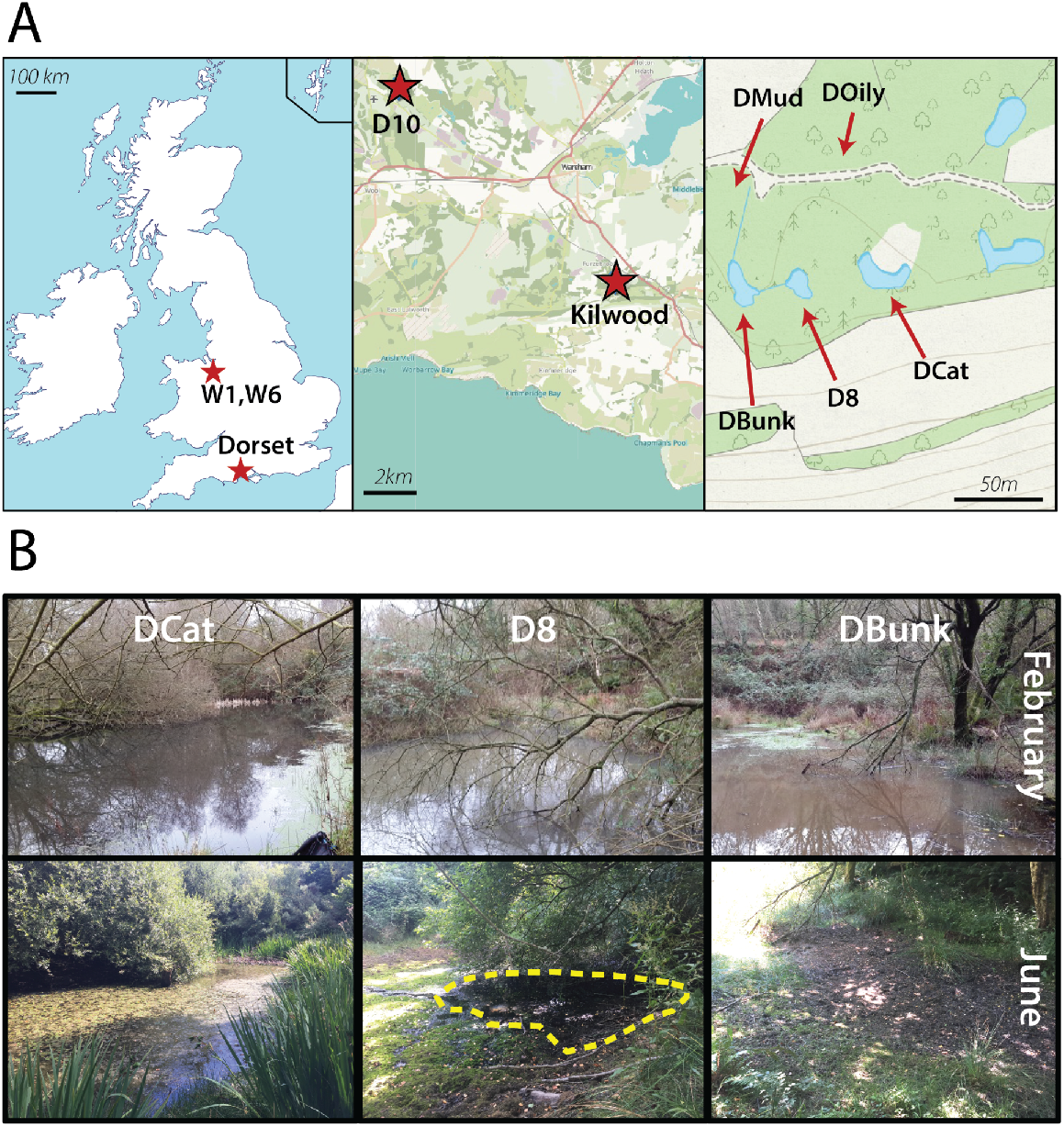
Location and features of the focal ponds. (A) Location of *D. pulex* sampling sites. The left panel shows the location of the focal populations, Dorset, and two distant ponds in Wales. The middle panel zooms in on the Dorset region and shows a neighboring pond, D10, along with the focal metapopulation at the Kilwood Coppice Nature Preserve, situated just north of the Purbeck hills. The right panel shows the location of the focal ponds DCat, D8, and DBunk, as well as two additional sampled ponds, DOily and DMud at the Kilwood Coppice (B) Pictures depicting water level in the three focal ponds in February and June, illustrating their variance in ephemerality. The outlined area in the June picture of D8 shows the borders of a remnant puddle. Map credits and references shown in the data accessibility statement. **<u>Supplemental Figure 1</u>. Quantification of relative water levels in D8 and DBunk.**

## Results

### Focal ponds vary in ecology

To characterize basic ecological aspects of the three focal ponds (DBunk, DCat, and D8, Figure 1A) we assessed water level over the course of the growing season and the presence of various *Daphnia* species. We documented that ponds varied in ephemerality: DBunk dried every summer, D8 periodically dried to a large puddle (e.g., during heatwaves in 2018 and 2019), and DCat exhibited relatively minor reductions in water level (Figure 1B). The geographically distant (11 km from focal ponds) Dorset pond, D10, also exhibited only minor changes in water level. The ponds rapidly filled in the fall after heavy rains (Figure S1). Thus, the growing season for these Daphnia populations is likely from fall into late spring or early summer. Multiple Cladocera species were found in the ponds: *D. pulex* was found in all ponds, *D. obtusa* only found in the most ephemeral pond, DBunk, and *Simocephalus* spp. found in the warmer spring months.

### Focal ponds vary in clonal diversity and mating dynamics

To characterize the clonal diversity of *D. pulex* from the focal ponds we sequenced >500 *D. pulex* genomes from samples collected across multiple years. These genomes consisted of 169 individuals fixed in the field and 344 lab-maintained isofemale lines (Table S1). Reads were aligned to a high-quality, chromosome-scale reference genome (L50=6, N50=10,449,493 bp, total assembly length=127,796,161 bp, BUSCO score=95%) derived from a *D. pulex* clone previously sampled from D8 (clone D8.4A, sampled in 2012). We assigned individuals to asexually related, clonal lineages based on pairwise identity by state (IBS, Figures 2A and S3A), and reconstructed a pedigree. We refer to clonal lineages that were sampled multiple times in the field as “superclones” ^42^ and individuals isolated from the field and propagated in the lab as “isofemale lines.”

**Figure 2.**
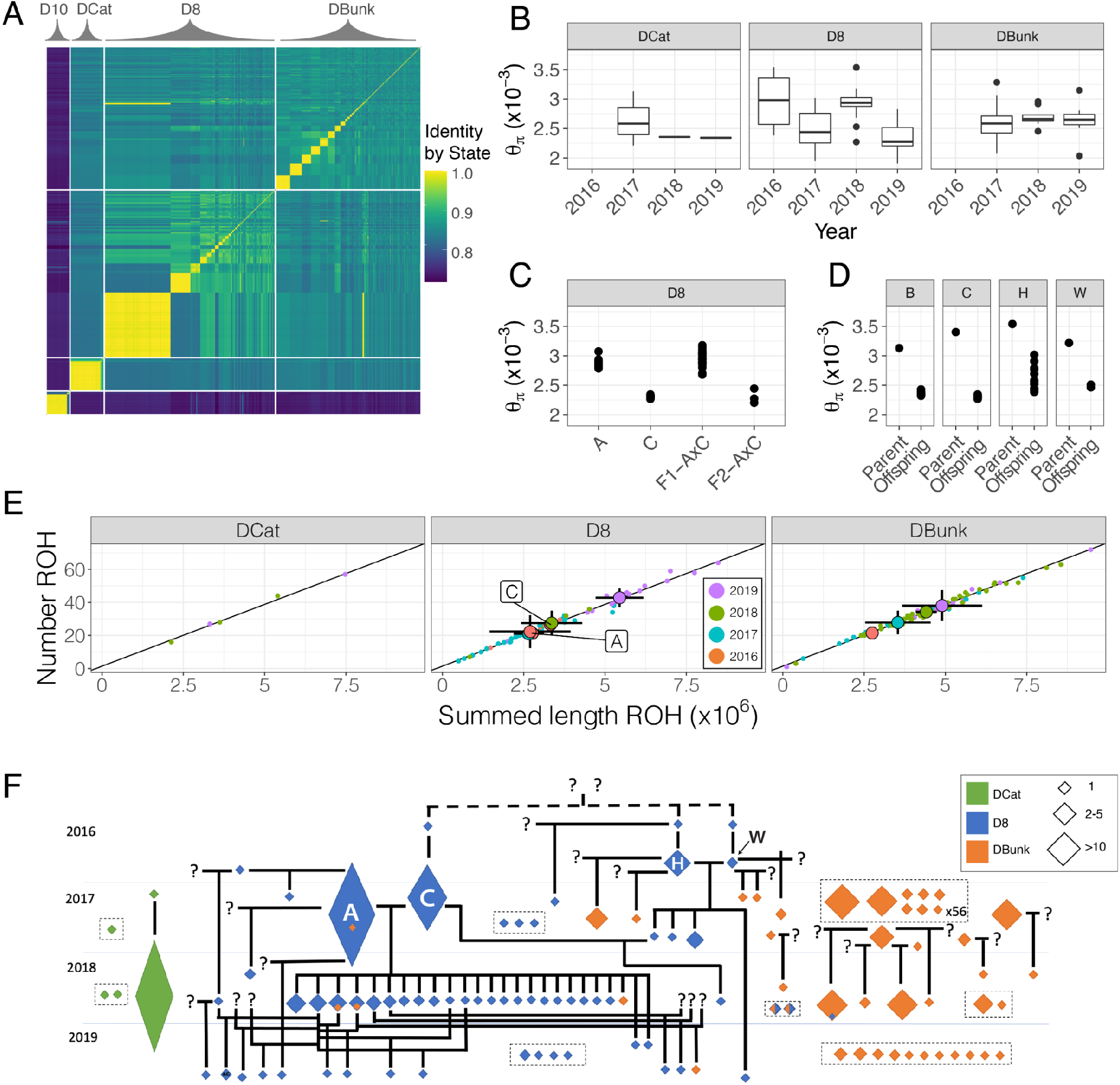
Clonal identity, genetic diversity, and pedigree. (A) Pairwise identity by state matrix generated using whole genome sequence data. Matrix includes 496 individuals from D10, DCat, D8, and DBunk. The largest clonal group identified in D8 is superclone A, the second largest is superclone C. (B) Estimates of genome-wide average heterozygosity across the three focal ponds. Each point represents the estimated heterozygosity for an individual clone. Diversity is stable through time in DCat and DBunk, but fluctuates in D8 due to cycles of inbreeding and outbreeding. (C) Genome-wide average heterozygosity between superclones A, C, their F1, and F2 progeny shows the direct consequence of outcrossing and intercrossing on diversity. (D) Genome-wide average heterozygosity between clonal lineages that selfed. (E) The number of runs of homozygosity (ROH) and total length of ROHs indicates that inbreeding and consanguinity are prevalent in these populations. Small points represent ROH estimates for each individual clone. Large points are yearly averages within each pond and error bars represent 95% confidence intervals. Individuals sampled in 2019 in D8 had significantly more, and longer ROHs than previous years (NROH: t_57_=3.92, *p*=0.00024; SROH: t_57_=3.99, *p*=0.00019). Yearly averages are only plotted for D8 and DBunk because there are too few unique clonal lineages in DCat. The diagonal black line is the estimate from a simple linear regression of nROH on sROH. (F) The inferred pedigree based on kinship and IBS_0_ (see Figure S2). The size of the diamond is proportional to the abundance of the superclone. **<u>Supplemental Figure 2</u>. Identification of clonal lineages and relationships**. **<u>Supplemental Figure 3</u>. Mutation accumulation in field sampled and lab reared superclones.** **<u>Supplemental Figure 4</u>. Introgression, population structure, and hybridization analysis of Daphnia samples**.

The relative abundance of clonal lineages varied across time and space and aligned with pond ephemerality (Figure 2A, 2E, and S4). The most permanent pond, DCat, was dominated by a single superclone that increased in frequency across all three sample years until it was the only clonal lineage sampled in 2019, leading to low clonal diversity across years (Shannon’s *H* range: 0-1.33, mean=0.51, Table S2). In contrast, the most ephemeral pond, DBunk, exhibited the highest clonal diversity (*H*: 2.04-3.59, mean=2.67) with many clonal lineages present across all time points. D8 was intermediate in clonal diversity (*H*: 1.02-3.05, mean=2.09), and exhibited the greatest fluctuations in diversity across years. Reduced clonal diversity was observed after an extended period of asexual reproduction (2017) when the pond likely did not go dry the previous summer, but higher levels of diversity were observed after the pond dried and sexual reproduction was enforced (2018, 2019). Persistence of clonal lineages across years also fit with observed patterns of ephemerality, with clonal lineages observed to persist across multiple years in both DCat and D8, but never in DBunk. In addition, patterns of mating varied with ephemerality. Consistent with reduced clonal diversity in more permanent ponds, selfing events and inbreeding between siblings were observed in the more permanent ponds (i.e., DCat and D8, but not DBunk; Figure 2E).

Patterns of genetic diversity within clonal lineages were consistent with extended periods of asexual reproduction. We found that new mutations within clonal lineages had an elevated nonsynonymous-to-synonymous ratio (*p*_N_/*p*_S_) relative to mutations in the same frequency class among outcrossing, sexually related clones (Figure S3). An elevated *p*_N_/*p*_S_ is expected during asexual reproduction as new mutations remain in the heterozygous state and are somewhat shielded from selection.

### Focal ponds do not vary in genetic diversity

To examine patterns of genetic diversity across space and time, we estimated average heterozygosity for each individual. Average pairwise heterozygosity (Θ_π_) among sampled lineages was ~0.0025 (Figure 2B), about one-third the magnitude of diversity estimates from North American populations ^43^. In contrast to clonal diversity and patterns of mating, genome-wide average heterozygosity did not consistently vary with ephemerality (Figure 2B), however diversity did fluctuate from year to year in D8. The fluctuations in D8 appear to be driven by this pond’s unique clonal dynamics, mirroring fluctuations in clonal diversity. Pairwise heterozygosity was high in 2016, then dropped in 2017 as two superclones came to dominate the pond. Mating between these two superclones led to an increase in pairwise heterozygosity in 2018, with inbreeding between the F1 hybrids leading to another reduction in pairwise heterozygosity in 2019. As expected, inbreeding and selfing events also led to reductions in genetic diversity (Figure 2C, 2D).

The recent pedigree structure of a population can have major impacts on the number and size of runs of homozygosity (ROHs) across the genome ^44^. In particular, populations that undergo bottlenecks or are consanguineous, as these Dorset populations appear to be (Figure 2F), are expected to have many, long runs of homozygosity (ROHs; > 0.1 Mb). Dorset individuals contained large ROHs (Figure 2E), although there was considerable variation among clonal lineages. Individuals sampled from DBunk had longer (t=2.57, *p*=0.011, Figure 2E) and more abundant ROHs (t=2.42, *p*=0.016, Figure 2E), relative to individuals sampled from D8. This observation is contrary to the simple expectation that D8 should have longer ROHs than DBunk, due to increased consanguinity ^44^, suggesting that the mechanisms influencing homozygosity in these populations are multifaceted.

### Coexistence of dominant clones was observed in the pond with intermediate ephemerality

One intriguing pattern in the spatio-temporal dynamics of clonal diversity was the existence of two superclones in D8 (hereafter referred to as superclones A and C, Figure 2D) that were relatively rare in 2016 (both frequency of 5%), came to dominate in 2017 (A:68% and C:22%), and then mated such that the majority of the D8 population in 2018 consisted of their F_1_ hybrids. Superclones A and C are themselves the product of sexual reproduction and are more distantly related than most individuals within the focal ponds in terms of kinship (A vs. C - 75 percentile) and (IBS_0_; A vs. C - 95 percentile; Figure S2E). Structure analysis ^45^ and tests of introgression ^46,47^ also support minimal gene flow from geographically distant *D. pulex* ponds, as well as two closely related *Daphnia* species, *D. pulex* and *D. obtusa* (Figure S4). Taken together, the results demonstrate that the divergence between superclones A and C is consistent with the segregation of loci present in the focal populations as a whole, with no need to invoke introgression from more distant *D. pulex* ponds or other *Daphnia* species.

### Coexisting dominant lineages exhibit a tradeoff between asexual and sexual reproduction

The fact that superclones A and C coexisted for some period of time (2016-2017), and then crossed to produce the majority of the population the following year (2018) suggests that these superclones may invest differently in sexual reproduction. A difference in reproductive investment is further supported by the lack of selfed A and selfed C genotypes among individuals sampled. A lack of selfed Cs in the field samples is particularly notable given that C was found to readily self when maintained in the lab (e.g. Figure 3E).

**Figure 3.**
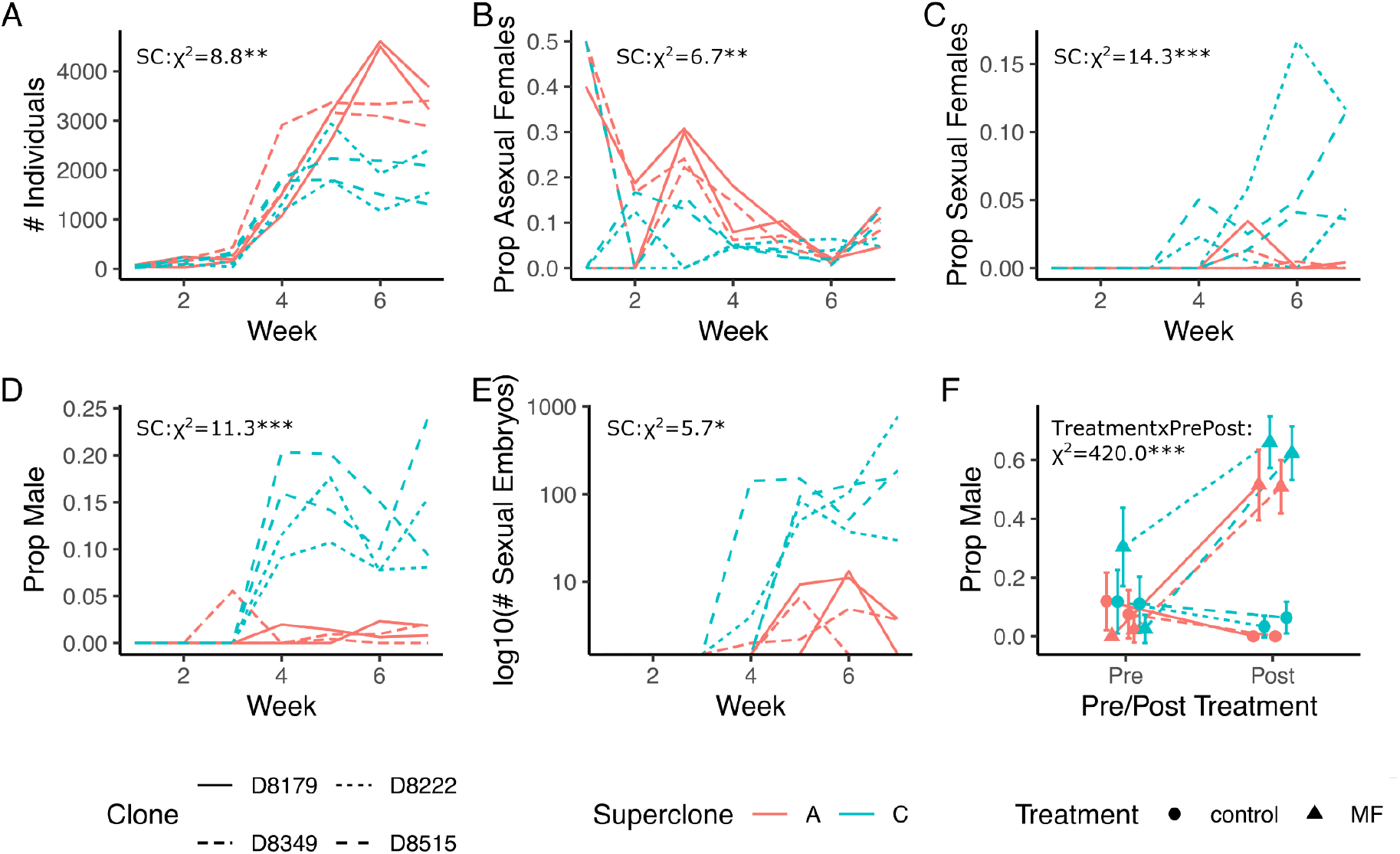
Superclone A and C invest differently in asexual and sexual reproduction. (A) Population size was higher for A isofemale lines than C isofemale lines when expanded in mesocosms, as was (B) the proportion of females reproducing asexually. (C) The proportion of females reproducing sexually was higher for C isofemales lines than A isofemale lines, as was (D) the proportion of males, and (E) the number of sexually produced embryos. (F) Exposure to methyl farnesoate (MF) led to increased male production in both A and C isofemale lines. Error bars represent 95% confidence intervals. Line types correspond to different isofemales lines. Two isofemales lines were used for each superclones (A: D8-179, D8-349; C: D8-222, D8-515).

To test the hypothesis that superclones A and C have differences in sexual investment, we established mesocosms inoculated with either superclone A or C isofemale lines. We tracked population size and demographic composition over seven weeks. Population trajectories in mesocosm expansion experiments demonstrated a clear tradeoff between asexual and sexual reproduction between superclones A and C. Superclone A isofemale lines achieved a higher total population size (Figure 3A, *χ*^2^=8.8, df=1, *p*=0.0030) and maintained a greater proportion of the population as asexually reproducing females for a longer period of time (Figure 3B, *χ*^2^=6.7, df=1, *p*=0.0096) than superclone C. In contrast, superclone C isofemale lines switched to sexual reproduction earlier, and to a greater extent, than superclone A females (Figure 3C, *χ*^2^=14.3, df=1, *p*=1.58×10^−4^). Perhaps most striking was the difference in male production. Superclone C isofemale lines produced substantially more males than superclone A and began doing so as soon as populations expanded between weeks 3 and 4 (Figure 3D, *χ*^2^=11.3, df=1, *p*=7.70×10^−4^). This difference in male production was maintained consistently throughout the rest of the experiment, and was replicable in multiple culture volumes (1L, *χ*^2^=14.7, df=1, *p*=1.27×10^−4^; Figure 4A; 250ml *χ*^2^=7.5, df=1, *p*=0.0063). Different allocation to sexual reproduction in terms of both male production and sexually reproducing females led to a marked difference between superclones in terms of sexual output, as measured by sexual embryo production (Figure 3E, *χ*^2^=5.7, df=1, *p*=0.017). Maximum weekly embryo production ranged from 84 to 770 for superclone C mesocosms, but only 5 to 13 for superclone A mesocosms.

**Figure 4.**
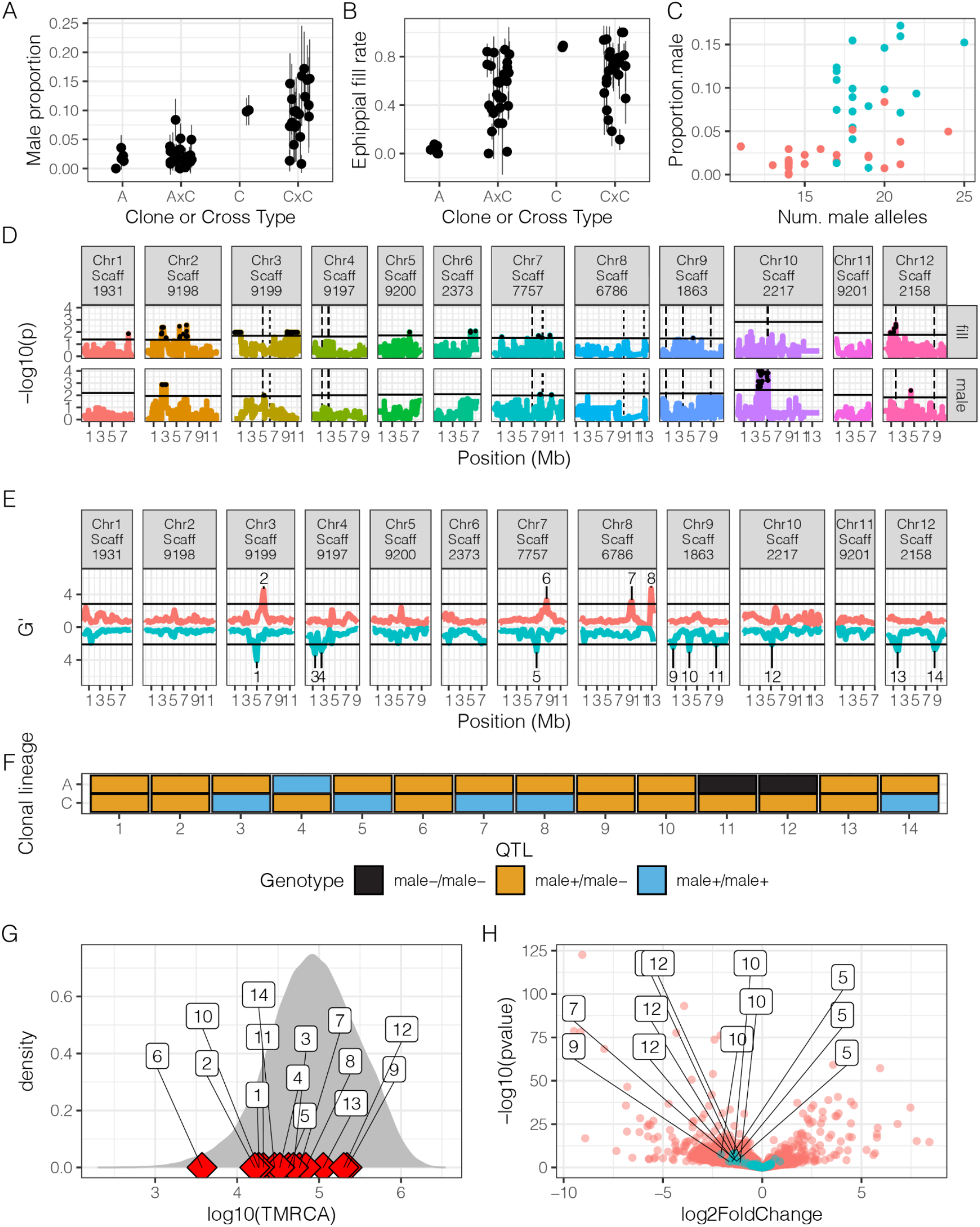
Genetic variation and mapping of variation in male production. Male production (A) rate and ephippial fill rate (B) in A, C, A×C F1s and C×C F1s suggests that sexual investment is polygenic. Points represent isofemale lines and vertical lines represent 95% confidence intervals (C) Male production rate is positively correlated with the number of QTL identified via Pool-Seq in both the A×C cross (red) and C×C cross (blue). Each point represents an isofemale line. See text for statistics on correlation between the number of *male^+^* alleles and male production rate. (D) QTL mapping in A×C F1 hybrids for ephippial fill rate and male production identified multiple peaks for each trait. Black points represent QTL regions that pass chromosome level permutation threshold. (E) Mapping of male production using pooled field samples identified multiple peaks, with some overlap with the A×C F1 hybrid mapping. To visualize overlap, the pooled sequencing peaks are plotted on the A×C F1 hybrid mapping figure as dashed lines. Horizontal line is the 5% FDR threshold. Values above the zero line represent PoolSeq replicate 1, and those below the zero line represent replicate 2. (F) Genotype for superclone A and C at each pooled sequencing QTL. (G) Time to most recent common ancestor (TMRCA; in *log_10_* transformed generations) for the 12 pooled sequencing QTL plotted against the genome-wide distribution. (H) RNAseq identified many genes differentially expressed between superclones A and C, some of which are located near QTL peaks. Blue points are genes within 50 kb of QTL identified in the PoolSeq, red points are all other genes, and genes that are near the PoolSeq peaks and are in the top 10% of differential expression genome-wide are labeled with their corresponding QTL number. For both (G) and (H), the number in the boxes correspond to the QTL number as identified in (E). **<u>Supplemental Figure 5</u>: Relationship between polar genotype and male production for F1 clones** **<u>Supplemental Figure 6</u>: Frequency of male polar genotype at each QTL in the three focal ponds**

### Unique mechanisms and origins of male limitation

Male production was the most pronounced phenotypic difference between superclones A and C in the mesocosm experiment. Male production in *Daphnia* can be triggered by exposure to methyl farnesoate, the innate juvenile hormone in *Daphnia* ^48^. However, previous work has shown that non-male producing clones in *D. pulex* ^28^ and *D. magna* ^30^ do not respond to methyl farnesoate, indicating that the loss of male production in these clones is associated with a loss of the ability to detect or respond to this chemical cue. To determine whether the difference in male production between superclones A and C was due to differing abilities to respond to methyl farnesoate, we exposed single A and C females to methyl farnesoate after their first clutch and tracked male production in subsequent clutches. The same four isofemale lines were used for the mesocosm experiment. Both A and C demonstrated a strong response to methyl farnesoate (40-60% male production rate) compared to controls (0-10%, Figure 3F, TreatmentxPre/Post: *χ*^2^=420.0, df=1, *p*<2.2×10^−16^). Superclone C females produced more males when exposed to methyl farnesoate than superclone A females (*p*=0.0079), consistent with C clones being higher male producers. Superclone A females also produced males when not exposed to methyl farnesoate, though at low frequencies (Figures 3C,F and 4A). Thus, superclone A is more accurately described as male-limited rather than non-male producing (*sensu* ^28,30^).

Previous work on variation in male production in North American *D. pulex* has shown that non-male production can be caused by adaptive introgression from the sister species, *D. pulicaria* ^28^. However, the *D. pulex* samples that we collected do not show signatures of recent hybridization (Figure S4C, S4D), and the fraction of the genome estimated to be derived from *D. pulicaria* or *D. obtusa* is low (<1%). In particular, superclones A and C show trivial amounts of ancestry with these two out-group species (~0.001%). Altogether, these findings indicate that the genetic architecture and evolutionary dynamics influencing variation in male production in the focal ponds are likely distinct from that observed in non-male producing clones in previous work.

### Crossing experiments indicate a polygenic architecture for male production

We examined aspects of the genetic architecture of male production by characterizing variation in reproductive investment of A×C (n=22) and C×C (n=20) *F_1_*s. Segregation of male production in A×C F1 hybrids suggested male production is consistent with a single, dominant male-limiting allele, as most A×C F1 hybrids exhibited male production rates similar to superclone A (Figure 4A). However, C×C F1s exhibited transgressive variation with male production rate (Figure 4B) suggesting that C is heterozygous for multiple loci affecting male production. These latter results argue against a single dominant male-limiting locus, in contrast to previous work ^27,28,30,38^.

To gain insight into the genetic architecture of male production, we sequenced the genomes of *F_1_* individuals and performed QTL mapping for male production rate and ephippial fill rate (number of sexual embryos per resting egg pouch, called an ephippium). Male production in the A×C cross identified multiple putative QTL, with the strongest association on chromosome 10 (Figure 4D). Ephippial fill rate amongst the A×C *F_1_* offspring also identified multiple QTLs with partial overlap to those identified for male production (Figure 4D).

### Mapping of male production using field samples further supports multiple QTL

We performed pooled sequencing of males and parthenogenetically reproducing females that were sampled and preserved in the field. We sequenced two pools of males (N=35 each) and two pools of parthenogenetically reproducing females (N=50 each) that were sampled from D8 in April of 2018. All males were made by parthenogenetically reproducing females and are therefore genetically identical to their male-producing mothers. Given that parthenogenetically reproducing females are likely to differ in their propensity for male production, regions that differ in allele frequency between the male and female pools represent candidate loci associated with male production.

Bulk-segregant analysis of the pooled sequencing data ^49,50^ identified 14 putative QTL associated with male production (Figure 4E). We used the identification and numbering of QTL in the pooled sequencing for the remainder of our analysis. Three of these QTL overlapped with peaks found for male production in the A×C F1 hybrid mapping (QTLs on chromosomes 2, 5, and 12), while one of the same QTL (chr. 2) plus one additional QTL (chr. 13) overlapped with peaks for ephippial fill rate. The overlap between the pooled sequencing and A×C *F_1_* hybrid mapping is only significant for ephippial fill rate (*p_perm_*=0.01).

Next, we asked whether the sign of the allelic effects at the 14 QTL identified via pooled sequencing is concordant with differences in male production between superclones A, C, and their *F_1_* offspring. If the QTL identified via pool-seq are enriched for true-positives, we expect a greater frequency of male producing alleles (*male*^+^) in superclone C, and a greater frequency of male limiting alleles (*male*^−^) in superclone A. Consistent with this prediction, we observe that superclone C contains slightly more *male*^+^ alleles than superclone A (*χ*^2^=4.86, df=2, *p*=0.087, Figure 4F). Notably, superclone C is homozygous at five putative QTL for the *male*^+^ allele, whereas superclone A is homozygous for the *male*^+^ at one QTL; superclone A is homozygous for the *male*^−^ allele at two QTL, whereas superclone C is not homozygous for the male limiting allele at any QTL.

We also examined the relationship between male production rate in the *F_1_*s and genotype, as polarized by the allelic effect inferred from the pool-seq QTL. Four of the 14 QTL had a significant relationship between genotype and male production at *p*<0.05, (two remained significant after Bonferroni correction), and of those four, three (including the two significant after correction) exhibited the expected relationship of greater male production in clones that were homozygous for the *male*^+^ genotype (Figure S5). We also observe a significant correlation between male production rate and the total number of *male^+^* alleles (Figure 4C) combined across both A×C and C×C crosses (binomial GLM: *χ*^2^=113.14, df=1, *p*=5×10^−26^) and within the A×C (*χ*^2^=18.87, df=1, *p*=1×10^−5^) or C×C cross (*χ*^2^=8.2, df=1, *p*=0.004).

### The ecological and evolutionary dynamics of male-production QTL

We hypothesized that the frequency of the *male^+^* alleles is correlated with the degree of ephemerality among ponds. We examined the allele frequency distribution of these QTL among ponds and calculated the expected frequency of the *male^+^* alleles by summing over the observed pond-level frequencies of each clonal lineage. In general, we do not observe a consistent relationship between *male^+^* allele frequency and the degree of pond ephemerality across all QTL (Figure S6), although there are some QTL which show striking patterns of differentiation consistent with our prediction (see below, *QTL_12_ is the best…).* These results are consistent with a model of adaptation at quantitative traits where differentiation is caused by stochastic allelic combinations across multiple loci ^51^.

We investigated the evolutionary history of the QTL associated with male production. We asked if alleles underlying these QTL are highly divergent, perhaps representing either recent introgression or old, balanced polymorphisms. Alternatively, young alleles may represent the product of rapid allelic turnover. We calculated the time to the most recent common ancestor (TMRCA), in generations, at every polymorphism genome-wide using the individuals from the Dorset ponds (Figure 4G). The 14 QTL identified via Pool-seq, in general, are younger than average but not exceptionally young. Only three QTL were older than the median genome-wide age, including the best candidate QTL on chromosome 10. These results suggest that QTL associated with male production are not highly divergent and allow us to reject the hypothesis that they represent introgression or stable balanced polymorphism. Rather, we suggest that male production is a trait that undergoes allelic turnover caused by recurrent selective sweeps.

### RNA-seq identifies candidate genes affecting male production

To assist in identification of candidate genes affecting male production between superclones A and C, we performed RNA-seq on whole, adult females. We first document abundant gene expression variation between the superclones (Figure 5). Principal component 1 clearly separates A and C (Figure S5A; t=-6.6, *p*=0.0022) and explains ~60% of the variation in gene-expression genome-wide. The magnitude of differential expression between these clones is similar to levels of differential expression previously documented between clonal lineages in other studies ^52,53^. There are 12 genes falling within 50Kb of five of the QTL peaks identified by PoolSeq that are differentially expressed (Figure 4H; Bonferroni corrected *p-*value < 0.05), and we consider these as candidate loci.

**Figure 5.**
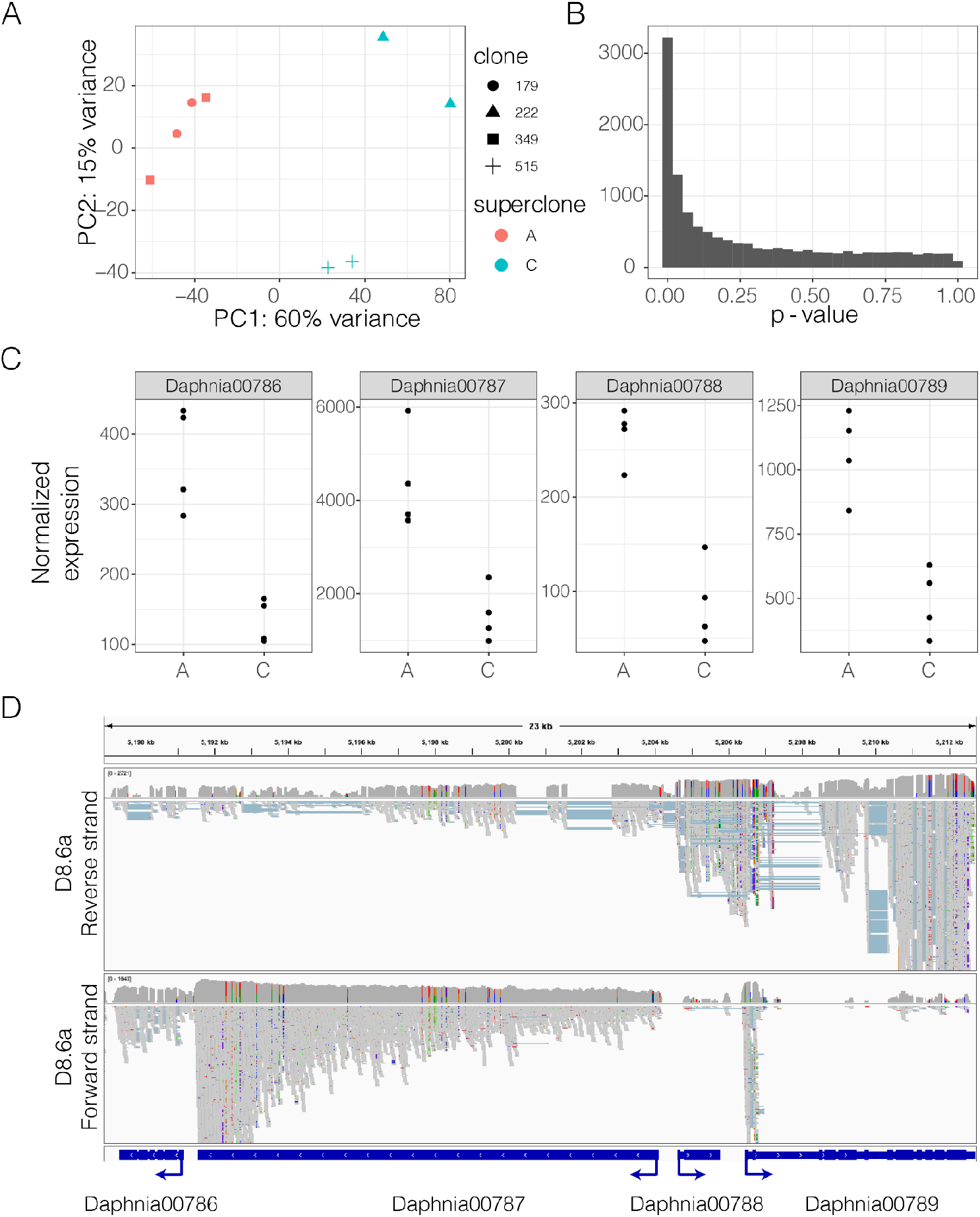
Differential expression between superclones A and C. (A) Principal component analysis shows that PC1 clearly separates superclone A and C, and explains 60% of the variance in gene expression. Clones within a superclone also clustered together. (B) The distribution of p-values contrasting differential expression between superclones. There are many differentially expressed genes. [C] shows the gene expression differences between superclones A and C for the 4 adjacent genes which are strongly differentially expressed. (D) IGView screenshot of the four-gene region from stranded RNA-seq libraries used for gene-model prediction. This screen-shot demonstrates that *Daphnia00787* is intronless, although there are spliced antisense transcripts possibly associated with *Daphnia00788* or *Daphnia00789*.

### QTL_12_ is the best candidate locus for male production

QTL_12_, located on chromosome 10 (Figure 4E), is of particular interest because of the confluence of multiple lines of evidence. QTL_12_ has the strongest signal in the A×C F1 hybrid mapping for male production (Figure 4D) and lines up with a QTL identified in the pooled sequencing mapping (Figure 4E). Superclone A is inferred to be homozygous for the *male*^−^ allele, and superclone C is inferred to be heterozygous *male*^+^/*male*^−^ (Figure 4F). Amongst the F1 A×C and C×C offspring, we find a strong association between genotype and male production that goes in the expected direction (binomial GLM: *χ*^2^=13.468, df=2, *p*=0.0011; Figures 6A and S5). QTL_12_ is strongly differentiated between DCat, D8, and DBunk in all three sampling years (Figure 6B). Notably, the frequency of the *male*^+^ allele among populations varies in the expected manner, with the highest frequency in the most ephemeral pond, DBunk, and the lowest frequency in the most permanent pond, DCat (Figure 6B). The *male*^−^ is closely related to the *male*^+^ allele (Figure 6C) and, consistent with our predictions, the *male*^−^ is derived.

**Figure 6.**
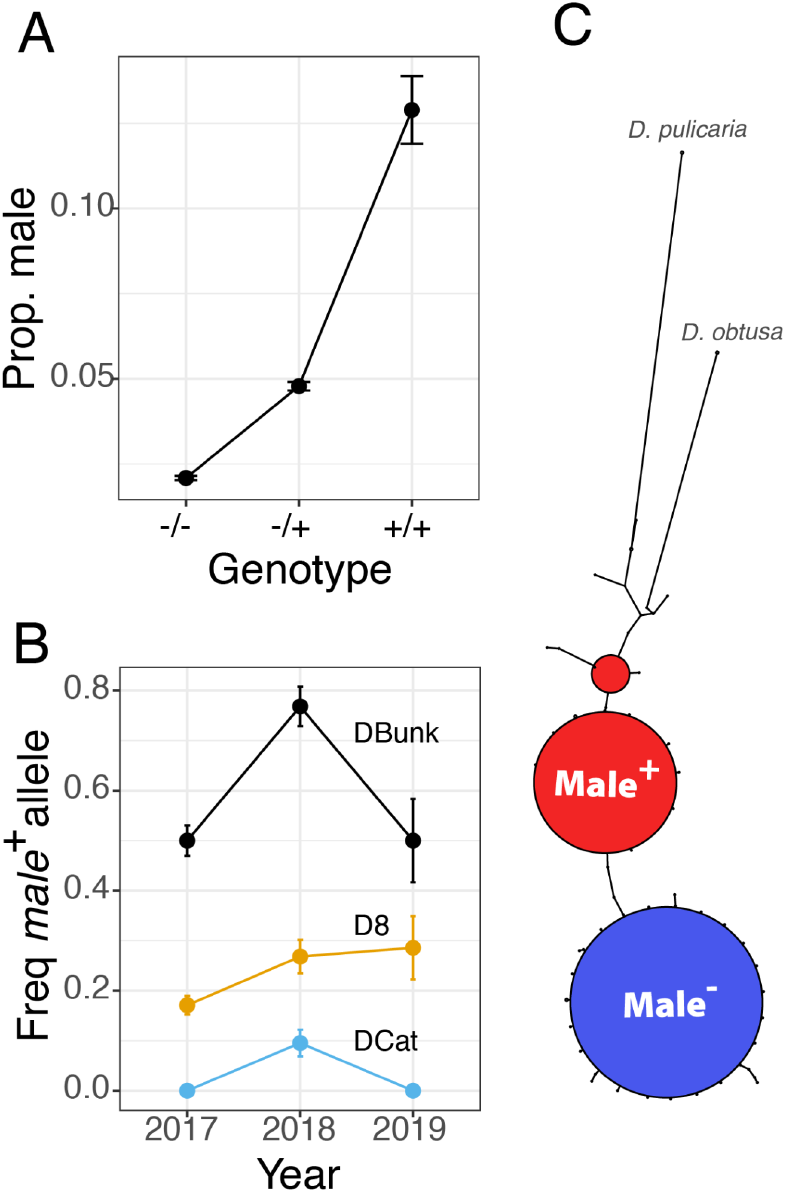
Attributes of QTL_12_. (A) Among the F1 offspring of the A×C and C×C cross, the proportion of male offspring increases with dosage of the *male*^+^ alleles, with the sign of allelic effect calculated from the Pool-Seq data. (B) QTL_12_ is strongly differentiated among ponds and the frequency of the *male*^+^ allele is aligned with the degree of ephemerality. (C) A haplotype spanning network plot shows that the *male*^−^ allele is derived from the *male*^+^ allele.

There are four genes close to the peak of QTL_12_ that are differentially expressed (Figures 4H and 5C). These four genes are amongst the top 7% of differentially expressed genes, genome-wide. These genes are immediately adjacent to each other (Figure 5D), and are consistently down-regulated in C compared to A (Figure 5C). It is likely that the correlated signal of differential expression among these four genes is a consequence of unannotated UTRs associated with one gene, *Daphnia00787,* overlapping with the open reading frames of the adjacent three genes (Figure 5D) and our use of unstranded RNA-seq of superclones A and C. *Daphnia00787* is the most plausible candidate gene of the four (see *Discussion*).

## Discussion

Herein, we studied the temporal dynamics of the genetic structure of *D. pulex* in a series of ponds that vary in ephemerality (Figure 1). Our work highlights the dynamic turnover of population structure through time in a facultative sexual and suggests that quantitative genetic variation in life-history traits can undergo rapid evolutionary turnover.

### Diversity and genetic structure through time

Clonal diversity, but not genetic diversity, varied in a pattern consistent with ephemerality (Figure 2). The more permanent ponds exhibited reduced clonal diversity, consistent with a longer asexual growing season and increased duration of clonal selection ^16,18^. Consistent with reduced clonal diversity, the more permanent ponds also exhibited a greater occurrence of inbreeding, including selfing and full-sib mating (Figure 2).

While individual selfing events reduced levels of heterozygosity, average heterozygosity was similar across all three ponds (Figure 2). There are several possible explanations for the consistently high levels of diversity despite the consanguineous pedigree that we observe. On the one hand, higher than expected levels of diversity could arise through neutral processes. For instance, neutral simulations that incorporated cyclical parthenogenesis showed that an excess of heterozygotes could be a consequence of drift early in the growing season ^54^. Diversity could also be maintained at high levels through migration and the subsequent reduction of inbreeding depression ^55^, although any migrant would face significant competition from the established *D. pulex* population ^16^. On the other hand, high levels of genetic diversity could be maintained via selective mechanisms. Haag and Ebert (2007) found that heterozygosity increased following clonal selection in an experimental *D. magna* system, potentially due to heterosis caused by associative overdominance ^56^. Fluctuations in environmental factors over the course of the growing season via micro-spatial variation in selection pressures or biotic interactions ^57,58^ could also contribute to the long-term persistence of variation.

Regardless of the mechanisms that have maintained genetic diversity, we provide clear evidence that clonal selection acts over short time periods. Although we do not have time-series data to directly observe clonal selection, our pedigree in D8 suggests that clonal selection occurred and that the rise in frequency of a limited number of clones was rapid. We were able to identify most of the clones involved in sexual reproduction in D8 — despite sampling only a small portion of the total population (21-117 out of hundreds of thousands) — indicating an effective population size that was orders of magnitude lower than the census population size. A striking example is that D8 in 2018 was dominated by F1 hybrids between only two superclones, suggesting that by the time sexual reproduction occurred in 2017, those two superclones had almost entirely taken over the population. Rapid clonal selection is consistent with observations from other *Daphnia* studies ^39,54,59–61^ and has also been observed in a variety of facultative sexuals ^18,19^.

### Genetic variation in sexual investment in a facultative parthenogen

Genetic variation in sexual investment has been observed in different *Daphnia* species as well as in other organisms capable of both sexual and asexual reproduction. The prevalence of genetic variation in sexual investment across multiple taxa suggests that male limitation arises frequently and independently and spreads rapidly in populations ^6,27,28,38,40^. In populations that are polymorphic for the potential to invest in sexual reproduction, male producing lineages tend to be rare ^41^, confirming the view that male production comes at a cost. We observe the male-producing superclone C to be at lower frequency in 2017 (22%) than superclone A (67%) the year prior to the hatching of A×C F1s (Figure 2). We do not observe any C×C offspring in 2018 despite ample selfing in experimental superclone C mesocosms in the lab, suggesting that superclone C may have been at an even lower frequency at the point in time ephippia were produced. The drying of D8 enforced sexual reproduction and the production of resting embryos and the following year, the population consisted almost entirely of A×C *F_1_* hybrids. This pattern demonstrates that although male producing clones may suffer a short-term fitness disadvantage, ultimately they can achieve high fitness if they are the primary source of males and dormancy remains linked to sexual reproduction. As a consequence, seasonal fluctuations in sexual reproduction can facilitate the maintenance of genetic polymorphism in sexual investment.

### Origin, stability, and potential mechanisms of male limitation

The evolutionary origin and genetic determinants of male limitation in the Dorset populations is also distinct from that of the North American system. The candidate gene underlying the major effect QTL identified in North American *D. pulex* ^28^ maps to chromosome 1 at positions 8,156,029-8,159,968 in the European *D. pulex* genome. This region is far from any putative QTL identified related to male production in the Dorset populations (Figure 4), supporting an independent evolutionary origin and mechanism. Moreover, we found no evidence of introgression or hybridization between the *D. pulex* populations that we study and English *D. pulicaria* or *D. obtusa* (Figure S4).

The best candidate gene that we identify could plausibly affect aspects of male-production. This candidate gene, *Daphnia00787*, is a large (4,179 amino acids), intronless gene (Figure 5D), orthologous to a toxin-like predicted toxin protein (*Tox-SGS*; Zhang et al., 2012) and a member of a gene family with orthologs only found in *Daphnia*, several blood-feeding insects and, notably, intracellular microbial symbionts including *Wolbachia, Cardinium,* and *Rickettsiella* ^63^. These cytoplasmically inherited symbionts influence sex ratios in a variety of arthropods ^64–66^ and it is therefore plausible that this toxin gene could be involved in sex-allocation in the *Daphnia* system that we identified here.

### Conclusion

We characterized the genetic structure of a *D. pulex* metapopulation across an ephemerality gradient and identified naturally segregating genetic variants affecting a key fitness trait, male production. It is likely that substantial genetic variation in other fitness related traits also segregates among these clones, and that such variation is maintained via selective processes. Due to the fact that the natural genetic structure of these populations resembles a multiparental QTL mapping panel, we envision that future work on the genetic basis of phenotypic variation in these, and other similarly structured populations, will yield valuable insight into the evolutionary forces that maintain genetic variation within populations.

## Materials and Methods

### Characterization and sampling of ponds

All field work and sampling were carried out in the Dorset region of the southern United Kingdom (Figure 1A). The focal metapopulation consisted of three geographically close (20-30m) small ponds located in the Kilwood Nature Reserve (50.642483, -2.091652) (Figure 1B,C). Two of these ponds, DBunk and D8 are intermittently connected, with DBunk downstream of D8; there are no obvious streambeds connecting DCat to either DBunk or D8. An additional pond, D10, located ~11 km away in the Higher Hyde Heath Nature Reserve (50.709379, -2.206421) was also part of this study and serves as a more distant population reference.

To characterize the local ecology and clonal dynamics of the focal populations we visited the four ponds at multiple time points across multiple years (2016-2019; Table S1). Water level of ponds was also noted and in 2019, time lapse cameras were installed at D8 and DBunk to monitor changes in water level. Tows were taken from each pond that still contained sufficient water to sample the *Daphnia* population. For the 2016 and 2017 time points, live individuals were shipped back to the lab and clonal lineages were established for later sequencing and phenotyping. For sampling points in 2018 and 2019, *Daphnia* were primarily fixed in ethanol in the field.

### Reference genome

Initial reference genome assembly was carried out using 10X Chromium sequencing. The *D. pulex* clone (D8.4A) used for the reference genome was sampled in 2012 from the D8 pond (Figure 1A) and maintained in the lab asexually since collection. Several hundred individuals from the clone were fed Sephadex G-25 superfine (cross-linked dextran gel) beads for 48 hours in order to clear their guts and minimize algal reads in downstream sequencing. After the 48 hours 70mg (wet weight) of *Daphnia* were flash frozen in liquid nitrogen and shipped to HudsonAlpha for high molecular weight DNA extraction, 10X Chromium library preparation, and whole genome sequencing on a single lane of HiSeqX. Assembly was carried out using Supernova-1.1.5 ^67^ (supernova mkfastq, supernova run, supernova mkoutput), with the input downsampled to 200 million reads and the output fasta made using the *pseudohap* style. Only resulting scaffolds over 1kb were kept for downstream analysis.

Scaffolding of the reference genome was achieved via Chicago and Hi-C sequencing (Dovetail Genomics, Scotts Valley, CA). Several hundred individuals from the same reference genome clone D8.4A were exposed to Sephadex G-25 beads and antibiotics (50mg/L each of streptomycin, tetracycline, and ampicillin) for 48 hours to minimize algal and bacterial contamination, and then flash frozen in liquid nitrogen. The resulting Chicago and Hi-C data was used to scaffold the 10X Chromium reference genome and resulted in an assembly with 9,202 scaffolds. Due to the fact that the *Daphnia* for the initial 10X Chromium assembly were not treated with antibiotics, many of these scaffolds are likely to be microbial in origin. By blasting the scaffolds against previously published North American *D. pulex* reference genomes (TCO and PA42) ^68,69^, 768 scaffolds were determined to be *Daphnia* in origin (positive hit to both genomes). Of these 768 scaffolds, 13 were over 50 KB and 12 were over 5MB. These 12 largest scaffolds were determined to correspond to the 12 chromosomes in the previously published *D. pulex* genomes (Table S3; ^68,69^).

To improve the reference genome assembly the reference genome clone was also sequenced on a Nanopore Minion flow cell. Around 100 individuals of the reference genome clone were exposed to antibiotics and Sephadex G-25 beads for 48 hours. DNA was extracted using Beckman-Coulter’s Agencourt DNAdvance kit. The library was constructed using Nanopore’s basic Ligation Sequencing Kit (SQK-LSK-109) and run on a single Minion SpotON flow cell (R9.4) for 48 hours using default parameters. Reads were basecalled using Albacore (version 2.3.3, Oxford Nanopore Technologies, Oxford, UK). The resulting MinIon reads (deposited at NCBI’s SRA: SRR14567272) were combined with the original 10X Chromium Illumina reads (filtered to only include reads that mapped to the good *Daphnia* contigs, thus excluding microbial reads) to make a hybrid genome assembly using MaSuRCA ^70^. This hybrid genome assembly along with the filtered 10X Chromium paired-end Illumina reads were then used to close gaps of Ns in the Dovetail HiC genome using GMCloser (v1.6.2, ^71^) with the -c option. The final genome size was 127,796,161 bp and 98.8% of that was contained in the 12 largest contigs. The genome sequence has been deposited at DDBJ/ENA/GenBank under the accession JAHCQT000000000.

Gene prediction was performed on the *D. pulex* genome using MAKER (v2.31.10; ^72^) in an iterative manner. First, *ab initio* gene prediction was performed by the programs SNAP (v2006-07-28; ^73^) and Augustus (v3.2.3; ^74,75^) using a multi-tissue *D. pulex* transcriptome (see *RNA-seq library preparation,* below) assembly obtained using StringTie with default parameters (v2.0; ^76^) and a protein database that combined the UniProt/Swiss-Prot and NCBI non-redundant database of reviewed arthropod proteins as well as the curated protein set for a related species, *D. magna* (http://arthropods.eugenes.org/EvidentialGene/daphnia/daphnia_magna_new/, accessed 5/6/2019). A total of three iterative runs of SNAP were used to refine the gene models. A final round of gene prediction was then performed using MAKER to produce the final gene set with an annotation edit distance (AED) cutoff of 0.5 ^72^. This gene set consists of 13,455 genes that encode 17,930 proteins. Genome annotation quality was assessed by BUSCO analysis ^77^ using the conserved core set of arthropod genes (~94.2% BUSCO score). Using the set of complete predicted *D. pulex* protein sequences, we used a functional annotation pipeline previously described ^78^ to assign putative functions to each predicted protein.

Repetitive elements were identified for masking in downstream analysis using RepeatModeler ^79^ and RepeatMasker ^80^. First, the RepeatModeler BuildDatabase option was used to construct a database from the D8.4A reference genome file. Next, RepeatModeler was run on the newly constructed database file. The resulting repeat library was then concatenated with the repeat family library constructed from the North American *D. pulex* genome, PA42 ^69^. RepeatMasker was run on the D8.4A reference genome using this combined repeat family library. Both RepeatModeler and RepeatMasker were run using the *ncbi engine* option.

We generated a list of regions that are masked from future analysis. First, we identified regions of the genome with very high or very low read depth, when mapping the Illumina reads generated for 10X linked-read sequencing back to the reference genome. We identified sites with the 1.5% lowest coverage and sites with 7.5% highest coverage sites and removed these sites from future analysis. In addition, we flagged stretches of Ns (±500bpo), the first and last 500 bp of sequence for each chromosome, and repetitive regions identified with RepeatMasker.

### Sequencing

Individual males, females, and pools of individuals were sequenced from samples collected across four years (Table S1). For the 2016 and 2017 samples, individuals were sampled from lines that had been established in the lab. For each of these lines, multiple individuals were exposed to antibiotics (streptomycin, tetracycline, and ampicillin, 50mg/L of each) and fed Sephadex G-25 beads. Five to 10 individuals from each line were used for DNA extraction (Agencourt DNAdvance kit, Beckman-Coulter). Individuals were homogenized using metal beads and a bead beater prior to DNA extraction. RNA was also removed using RNase followed by an additional bead cleanup. DNA was quantified using the broad-range Quant-iT dsDNA kit (ThermoFisher Scientific) and normalized to 1 or 2 ng/ul prior to library construction. Full genome libraries were constructed using a scaled down Nextera protocol ^81^. Libraries were size-selected for fragments ranging from 450-550bp using a Blue Pippin and quality checked using a BioAnalyzer. For 2018 and 2019 samples, For the 2018/2019 samples, DNA was extracted from single individuals (females with parthenogenetic eggs or males) fixed in ethanol in the field. Normalization and library construction were carried out in the same way as for 2016 and 2017 samples, except the samples were not RNAse treated due to low DNA concentration.

Pools of males and females were made up of individuals sampled from D8 on 04/29/2018 and fixed in ethanol in the field. There were two pools of 50 females and two pools of 35 males. Libraries were constructed using NEB’s NEBNext Ultra II DNA Library Prep Kit for Illumina.

Three additional individual *D. pulex* from two more northern populations in the UK (W1 and W6), 15 *D. obtusa* (14 from DBunk, and one from a nearby pond also in the Kilwood Nature Reserve, DBarb), and 5 *D. pulicaria* sampled from a single population in the UK (53.581442, −0.312748) were also sequenced to serve as geographically and taxonomically distant references.

Samples from 2016 (N=52) were sequenced on two lanes of HiSeqX. Samples from 2017 were sequenced in three batches (N=94, 94, and 190; 3, 3, and 6 lanes of HiSeqX respectively), with some samples repeated between batches if initial read depth from early sequencing runs was too low. Initial samples from 2018 (N=94) were run on a single lane of NovaSeq 6000 S4 300. Samples from 2019 (N=61), additional samples from 2018 (N=33), and the male and female pooled samples from 2018 were pooled and sequenced on a single lane of NovaSeq 6000 S4 300. Fastq samples are available on NCBI’s SRA (PRJNA725506). See Table S1 for accession numbers for individual samples. Pooled samples: SAMN19227056-SAMN19227059.

### Mapping, SNP calling, and annotation

For each of the individual samples, Nextera adaptor sequences were removed using *Trimmomatic* v0.36 ^82^. Next, overlapping reads were merged using *PEAR* v0.9.11 ^83^. Assembled and unassembled reads were separately mapped to the European *D. pulex* genome using *bwa mem* ^84^. The entire reference genome was used for mapping, but only reads that mapped to *Daphnia* scaffolds were retained for further analysis. Reads that were primary alignments and had mapping quality scores greater than 20 were output into bam files. For samples that were sequenced across multiple lanes, samtools ^85^ was used to merge bam files from each lane, and then PCR duplicates were removed using *MarkDuplicates* function of *Picard* ^86^. GATK (v4.0; ^87,89^) was used to call SNPs. gVCFs for each sample were made using GATK’s HaplotypeCaller tool, and then a single VCF was made using GATK’s GenotypeGVCF tool. We performed functional annotation of the VCF file using *SnpEff* (v4.3t, ^88^) utilizing the gene predictions described above.

### SNP filtering

First, we filtered out SNPs that were within 10 base pairs of indels; indels we also removed from analysis. SNPs were hard filtered using GATK’s recommendations for organisms with no reference SNP panel. Specifically, the following filters were first set using GATK’s VariantFiltration tool (QD < 2, FS > 60, MQ < 40, MQRankSum < −12.5, and ReadPosRankSum < −8), and SNPs that did not pass this filter were removed using GATK’s SelectVariants tool. Individual genotype calls with low quality scores (GQ < 10) were then set as missing data. The resulting VCF consisted of 3,719,919 SNPs.

SNPs were removed if they fell in regions that were flagged as being near stretches of Ns or the ends of chromosomes, as well as in areas of high or low read depth (described above in the reference genome section). Together, this filtration resulted in a loss of 651,900 SNPS, leaving 3,068,019.

SNPs were filtered from regions flagged by Repeat Masker (described above in the *Reference Genome* section). Filtering SNPs from flagged repeat regions resulted in a loss of an additional 320,003 SNPs. Triallelic SNPs were also removed, resulting in a further loss of 81,624 SNPs. SNPs were then filtered based on read depth across all samples, with the bottom and top 5% of SNPs being dropped, which resulted in a loss of 266,515 SNPs, leaving 2,399,653 SNPs. This set of filtered SNPs included SNPs polymorphic within *D. pulex*, as well as SNPs that are fixed within *D. pulex*, but are either different between *D. pulex* and one of the outgroups, or polymorphic within one of the outgroups. This SNP set will be referred to as the “total filtered SNP set”.

For analyses restricted to *D. pulex* samples, a second filtered SNPs set was used. For this SNP set, only SNPs that were polymorphic within the *D. pulex* samples and had been genotyped in at least half the *D. pulex* samples were retained. This SNP set, consisting of 510,805 SNPs, will be referred to as the “variable *pulex* SNP set”. Finally, this variable *pulex* SNP set was also LD pruned using the snpgdsLDpruning function in SNPRelate ^90^, with a minor allele frequency cutoff of 0.001, a missing rate of 0.15, a maximum window size of 500 bp, and an LD threshold of 0.1. This LD pruned SNP set had 150,455 SNPs and is referred to as the “LD pruned, variable *pulex* SNP set”.

### Clonal assignment

Individual clones were assigned to clonal lineages using genome wide estimates of identity by state (IBS) calculated using the *snpgdsIBS* function in *SNPRelate* ^90,91^, excluding singleton SNPs (1/2N) and SNPs with a missing rate greater than 15%, and using the LD pruned, variable *pulex* SNP set. Based on the distribution of pairwise IBS (Figure S2), an initial cutoff of 0.965 IBS was chosen as a threshold, above which two individuals are considered to belong to the same clonal lineage (asexually related to one another). Note that these IBS values are calculated based on the table of polymorphic sites, and not the whole genome and thus the denominator for these statistics is the number of polymorphisms and not the genome size. Assuming a genome size of 120Mb and that the number of polymorphisms used for this analysis is ~500,000, maximum divergence (Dxy) between individuals of the same superclone is 1.44×10^−4^.

### Genealogical relationships between isolates

To further investigate patterns of relatedness, we calculated *IBS_0_* and kinship coefficients using the program KING ^92^. KING was run using the “kinship” command on the variable *pulex* SNP set (non-LD pruned), with the input data filtered to include SNPs with a minor allele frequency cutoff of 0.05. The distribution of *IBS_0_* and Kinship mostly confirmed the clonal lineage assignment based on the IBS threshold. However, there were three individuals that appeared as outliers within their clonal lineages in terms of Kinship. Specifically, the Kinship values were lower than that expected for clonal relationships, but higher than expected for outcrossing parent-offspring relationships (e.g. Figure S2C,D). In one case, based on patterns of SNP segregation, the outlier individual was identified as the parent of the lineage, with the lineage resulting from a selfing event. Interestingly, in three other examples of selfing, the chosen IBS threshold did separate the parent from the offspring lineage. In two other cases, the relationship was not as clear, and the outlier individuals were classified instead as close relatives of their respective clonal lineages. In all three cases, the individuals were removed from their respective clonal lineages.

The distribution of pairwise *IBS_0_* and Kinship calculated in KING were also used to identify parent-offspring relationships and construct a pedigree. Similar to clonal relationships, parent-offspring pairwise comparisons (whether due to selfing or outcrossing events) are expected to have an *IBS_0_* of roughly 0, with Kinship values being highest for clonal comparisons, followed by selfed parent-offspring, and then outcrossed parent-offspring. Graphing *IBS_0_* and Kinship resulted in clear clustering of these three types of relationships (Figure S2B), which were used to identify parent-offspring relationships and construct a pedigree for the three focal ponds across sample years. Selfed parent-offspring relationships were confirmed by checking segregation patterns of parental heterozygous sites in the offspring. Segregation patterns of parental SNPs were also used to check outcrossed parent-offspring relationships where possible. For example, A×C F1 hybrids were confirmed to be heterozygous for all SNPs that are fixed differences between A and C.

### Diversity estimates and runs of homozygosity

ROHan v1.0 ^93^ was used to estimate pairwise genetic diversity for each individual (Figure 2) and to identify runs of homozygosity. For this analysis we used the 12 major chromosomes and a transition/transversion ratio of 0.81 ^94^.

### Mutation accumulation

To study patterns of mutation accumulation in the wild, we identified 8 wild-caught individuals from 4 superclones (range of individuals per superclone = 9-127). We randomly selected 8 individuals per superclone and estimated *p*_N_/*p*_S_ for new mutations (those segregating at less than 25%) and for shared variants (those segregating at ~50%). We generated confidence intervals on pn/ps by bootstrap resampling (n=100).

### Phasing and imputation of wild-caught individuals

We identified one representative per superclone among the 169 wild-caught *D. pulex* (Table S1) by selecting the clone with the highest read depth. We performed read-backed phasing using *whatshap* (v1.1, ^95^), and performed population based phasing and imputation using *shapeit* (v4, ^96^). We generated consensus genotypes for two outgroups - *D. pulicaria*, *D. obtusa*, at sites that were polymorphic in the phased-imputed dataset.

### Hybridization and Introgression analysis

We tested for potential hybridization and introgression between *D. pulex* and its outgroups in two ways. First, we used *Dsuite* ^46^ to calculate *f_4_* and *f_4_*-ratio statistics ^47^ using the phased, imputed dataset. Next, we used the *snmf* function in *LEA* (v3.12, ^45)^ to estimate individual admixture components using the phased, imputed dataset. For the *snmf* analysis, we varied K from 1-20 and performed 30 replicate runs. We selected the value of K with the lowest cross-entropy score as the optimal number of clusters (k=8).

### Phenotypic differentiation between A & C

To test for phenotypic differences between the two dominant superclones in D8 (A and C), two field collected (2017) isofemale lines of each superclone were expanded in mesocosms (A: D8-179, D8-349, C: D8-222, D8-515). We tracked population dynamics over the course of eight weeks. First, each isofemale line was expanded in two tanks containing 15 liters of artificial hard water (ASTM; ^97^) with seaweed extract (marinure; Wilfrid Smith Std., Northans, UK) in a Percival incubator set at 18 degree C under long days (16L:8D). 60 females just prior to first reproduction were isolated from the mesocosms once in the expansion phase, and used to establish the experimental mesocosms.

Eight experimental mesocosms were established, two for each isofemale line (2 field isolates × 2 clonal lineages × 2 replicates = 8 tanks). Each experimental mesocosm consisted of a fish tank containing 15 liters of ASTM with marinure placed in the same Percival incubator set at 18 degree C under long days (16L:8D). For the first five weeks each tank was fed 95,000 cells/mL *Chlorella vulgaris* M/W/F, and then were switched to 142,000 cells/mL algae for weeks 6-7. Tanks were checked weekly after initial establishment for seven weeks. To check the tanks, each tank was stirred up well to evenly suspend all individuals, and then a liter of media was removed from each tank and sieved to isolate the *Daphnia*. These *Daphnia* were fixed in ethanol and later sorted into the following demographic classes: females with parthenogenic eggs, females with ephippia, females with neither parthenogenic eggs or ephippia, female neonates, juvenile males, and adult males.

At each sample time point, all loose ephippia were also removed from each tank. All ephippia were counted and a subset were dissected to determine fill rate. These fill rates were used to estimate total sexual embryo production (estimated fill rate × total number of ephippia = estimated sexual embryo production). Total population size was calculated as the total number of individuals in the sample times 15.

Significant differences between superclone for total population size, proportion asexual females, proportion sexual females, male production, and sexual embryo production were tested using lmer or glmer in R (lme4; ^98^. Differences in population size and sexual embryo production were tested using lmer (gaussian error model), with isofemale line and week included as random effects, and comparing the fit of the model with and without the inclusion of superclone as a fixed effect. Anova was used to test for a significant improvement in fit when including superclone. Proportion asexual females, sexual females, and males were analyzed in a similar fashion, but using a binomial error model

The same four isofemale lines used for the mesocosm experiments (A: D8-179, D8-349, C: D8-222, D8-515) were exposed to methyl farnesoate to determine their response in regard to male production. Individual neonates released within 24 hours were placed in 50 ml jars in ASTM with marinure. The second clutch neonates from these individuals were used for the experiment. Individual neonates were placed in 50 ml jars in ASTM with marinure and checked M/W/F. When the first clutch was observed, the female was moved to a new jar and the neonates were counted and scored as male or female. Females in the methyl farnesoate treatment were placed in jars containing 50 ml of ASTM plus marinure and methyl farnesoate at a concentration of 400 nM. Females in the control treatment were placed in jars containing 50 ml of ASTM plus marinure and ethanol to control for any side effects of the ethanol in which the methyl farnesoate was resuspended. Jars were media changed M/W/F and checked for neonates, which were counted and scored as male or female. Jars were followed for 4-6 clutches, although in a few cases individuals died prior to the fourth clutch. Altogether 83 neonates were included in the experiment (N=7-14 per trmt/field isolate combination), 71 of which survived to at least the third clutch (5-11 per trmt/field isolate combination). Throughout the experiment jars were fed 200,000 cells mL^−1^ *Chlorella vulgaris* M/W/F and were maintained under 16:8 light:dark conditions at 20°C.

For analysis, the proportion of male neonates from the first clutch was used as a pre-treatment estimate of male production. The summed proportion of males from clutches 3 and higher were used for post-treatment estimates of male production. Clutch 2 was not included in the analysis as this clutch was not evenly exposed to methyl farnesoate across jars. Since jars were checked M/W/F and females were moved into treatment after the first clutch was observed, the exposure of the second clutch to methyl farnesoate was variable, but the third and higher clutches were fully exposed for all females. A significant effect of methyl farnesoate exposure was tested using a generalized linear model, with pre/post treatment exposure, treatment (methyl farnesoate vs control) and clonal lineage (A vs C) as main effects as well as all two-way interactions. A binomial distribution was used, and the total number of neonates was used as a weighting factor.

### Generation and phenotyping of F1 A×C and C×C offspring

Patterns of segregation in ephippial fill rate and male production were examined in both selfed C and A×C F1 hybrid clonal lineages. Selfed C and A×C F1 clonal lineages were obtained from the mesocosms, described above. Ephippia isolated from the mesocosms were hatched and clonal lineages were maintained in the lab for several months prior to the experiment. Selfed C lineages (N=24) came from C tanks as well as A/C mixed tanks. A×C F1 hybrid lineages (N=26) came either from A/C mixed mesocosms or were isolated from the field. No selfed A or selfed C clonal lineages were isolated from the field. The four A/C superclone field isofemale lines (A: D8-179, D8-349; B: D8-222, D8-515) were also included. One liter jars were established for each clonal lineage and maintained at high densities (average of 50 reproductively mature females, 125 individuals total) and maintained for multiple weeks. The jars contained ASTM plus marinure, were fed 200,000 cells mL^−1^ *Chlorella vulgaris* M/W/F, and were maintained under 16:8h light:dark conditions at 20°C. The artificial hard water was changed via pouring over every two weeks (leaving a small amount of original water in the jar), maintaining high population sizes, and all loose ephippia were collected, counted, and dissected to score fill rate. Most clonal lineages were run in two replicates, while a few had either one or three replicates, and the A/C isofemales lines had five replicates. Jars were checked between 2 and 16 times, with a mean of 6. At the end of the experiment, the jars were sieved, and the *Daphnia* were fixed in ethanol. *Daphnia* were sorted into demographic classes and counted, and proportion male was calculated.

### Phasing and imputation of A×C and C×C F1 offspring

We performed transmission based phasing on 50 unique lab- and field-generated F1s using *rabbit* (v3.2, ^99^). All field-caught individuals (n=10) were inferred as AxC F1s based on patterns of segregation described above (see *Genealogical relationships…*). There were 16 lab-generated A×C and 24 lab-generated C×C genomes sequenced. First, we built consensus genotypes for the parental genomes (A and C) and selected the 5000 informative markers between these strains. Next, we used the *magicImpute* function to impute missing data in the offspring and to phase the parental genomes. Finally, we used *magicReconstruct* to generate phased genomes of the recombinant offspring using the Viterbi decoding algorithm.

### QTL mapping using F1s

Using the phased and imputed genomes of recombinant individuals, we identified the 5436 unique polymorphisms. These SNPs represent “tag” SNPs and are in perfect linkage with the remaining 115399 informative markers identified between A and C. We used this set of markers for QTL analysis, and later propagated the signals of association at the tag SNPs to the entirety of the linkage block.

We performed QTL mapping using *lme4qtl* ^100^ and generated the additive genetic relatedness matrix (GRM) using the *A.mat* function in *rrblup* (v4.6.1, ^101)^. We generated a new GRM for each of the SNPs we tested in order to avoid proximal contamination ^102^. We performed 100 permutations to generate an empirical false discovery rate ^103^, which we calculated for each chromosome separately.

### Bioinformatics and analysis of Pool-Seq samples

Pool-seq samples of males and females bearing parthenogenic offspring were generated, as described above. We first estimated allele frequencies in the Pool-Seq samples using the ASEReadCounter function of GATK ^89^ at the ~500,000 SNPs that are part of the “variable *pulex* SNP set”.

Previous sequencing of 2018 D8 clones in the lab had indicated that in 2018 the D8 population predominantly consisted of AxC F1 hybrids (Figure 2E). We confirmed that the Pool-seq samples were primarily F1 hybrids between A and C in two ways. First, we calculated the proportion of sites segregating in the Pool-Seq data that are also polymorphic between A and C. Superclones A and C are different genotypes at 98% of sites with MAF > 5% in the Pool-Seq data. Second, we partitioned allele frequencies in the Pool-Seq samples based on the nine F1 genotypic combinations between A and C and found that observed allele frequencies were close to the expected value. Because our Pool-Seq samples are composed of genetically diverse F1s we are justified in performing bulk-segregant analysis as typically applied to experimental crosses.

We used the *G’* test ^49^ as implemented in *QTLSeqR* ^104^ to perform bulk-segregant analysis. First, we took the reference and alternate allele counts calculated by *ASEReadCounter* and down sampled them proportional to the effective read depth ^105,106^, which factors out the double binomial sampling that occurs during Pool-Seq. We calculated the *G’* statistic filtering for sites with MAF > 15%, minimum total read depth of 20, minimum sample depth of 20, and a window size of 250,000bp. We calculated *G’* pairing one of each replicate pool of females and males. We corrected for multiple testing using the Benjamini-Hochberg FDR method ^107^.

### RNA-seq library preparation

We conducted RNA sequencing on two A clones and two C clones, using two biological replicates of each. We snap-froze 20 females for each biological replicate in liquid nitrogen and stored samples at −80 °C. We extracted RNA using the RNAdvance Tissue kit (Agencourt) following manufacturer’s instructions, including the optional DNase treatment step. RNA integrity was verified using an Agilent Bioanalyzer (all RIN > 6.2). We isolated poly-adenylated RNA from 1 ug of total RNA using the NEBNext Poly(A) mRNA Magnetic Isolation Module (NEB #E7490) and constructed sequencing libraries with the NEBNext Ultra II RNA Library Prep kit (NEB #E7770) using dual indices (NEB #E7600). We quantified libraries by Bioanalyzer and with a Quant-iT kit (Thermo Fisher) and pooled the eight libraries in equimolar concentrations. The pooled libraries were sequenced in one lane of Illumina HiSeq X with 150 bp paired-end reads. The resulting fastq files were deposited in NCBI’s SRA: SRR14572418-SRR14572425.

We conducted RNA sequencing a *D. pulex* clone (D8.6A) that was collected from the same pond as the reference genome (D8) in 2012 and maintained as iso-female lineages in artificial hard water (ASTM) with seaweed extract (marinure) under standard conditions in the lab. Prior to RNA extraction, this clone was maintained under standard conditions for three generations and fed daily. Animals were sieved, rinsed with ASTM and snap-frozen in liquid nitrogen at late embryonic stage (E4; 286 pooled embryos per sample), and early and late first instar (I1.1early, I1.1late; 143 pooled animals per sample each). For sample preparation, 896 μl diluted methanol was added to each sample, and samples were homogenized for 2 × 10s bursts at 6400 rpm (Precellys). Samples were placed on dry ice and 300 μl sample was transferred into an RNAse free tube and flash frozen in liquid nitrogen. RNA from samples was subsequently extracted using the RNAdvance Tissue kit (Agencourt) following manufacturer’s instructions, including the optional DNase treatment step. RNA integrity was verified using an Agilent Tapestation 2200 with High Sensitivity RNA screentapes (all RIN > 6.2). RNA libraries were produced using the Biomek FxP (Beckman Coulter A31842) with NEBNext Ultra II Directional RNA Library Prep Kit (New England Biolab E7420L) and NEBnext Multiplex Oligos for Illumina Dual Index Primers (New England Biolabs E7600S), using provided protocols and 500 ng of total RNA. Constructed libraries were assessed for quality using the Tapestation 2200 with High Sensitivity D1000 DNA screentape. Multiplex library clustering and sequencing was performed upon the HiSeq2500 (4 lanes rapid run 2x 100bp) by BGI Copenhagen. Fastq files are available at NCBI’s SRA (PRJNA727995).

### RNA-seq bioinformatics

RNA-seq reads were mapped to the reference genome using STAR (v2.7.2b, ^108)^ using the following parameters: --outFilterMatchNmin 0 -- outSJfilterReads Unique --outSJfilterCountUniqueMin 20 1 1 1 --alignIntronMax 25000 -- outFilterMismatchNmax 20 --outFilterIntronStrands RemoveInconsistentStrands -- sjdbOverhang 100. Quality control of mapped reads and post-mapping processing was performed using QoRTS (v1.3.6, ^109)^. Differential expression analysis between superclones A and C was performed using DESeq2 ^110^.

### Haplotype network analysis

We estimated a haplotype network by using whole haplotype sequences extracted from the phased VCF containing one representative per superclone and including sequences from outgroup taxa. Diploid haplotypes for the *Daphnia00787* gene were extracted using *bcftools* ^85^ (consensus function) and the reference genome of *D. pulex* sliced to the region corresponding to Scaffold “2217_HRSCAF_2652” from 5,191,562 bp to 5,204,101 bp. Before analysis, haplotype sequences were reversed and complemented using the string unix functions *rev* and *tr* ACGT TGCA, respectively. Individual haplotypes were concatenated into a single FASTA file and loaded into R using functions from *ape* (Paradis et al., 2004). Network analysis was done using the package *pegas* ^111^.

### Allele age analysis

Estimates of allele age (TMRCA) were done using the GEVA program ^112^ and using the following population genetic parameters: recombination rate = 1.60e-8, mutation rate = 5.69e-09 and effective population size = 862,000. GEVA was implemented on the phased VCF containing one representative per superclone excluding all outgroups. Prior to analysis, we filtered SNPs with mAF < 0.01.

### *Daphnia00787* orthology

Orthology of *Daphnia00787* was assessed by blasting the reference sequence of the gene against the NCBI database using the *blastp* algorithm with default settings.

### Statistical analysis and plotting

Statistical analysis was performed using R version 3.5 to 4.0.5 ^113^. The following packages were used for general analysis and plotting: *ggplot2* ^114^, *cowplot* ^115^, *patchwork* ^116^*, viridis* ^117^, *leaflet* ^118^, *data*.*table* ^119^, *foreach* ^120^, *doMC* ^121^, *SeqArray* ^91^.

## Author contributions

<u>Karen Barnard-Kubow</u>: Conceptualization, Data curation, Formal analysis, Funding acquisition, Investigation, Methodology, Resources, Software, Visualization, Writing - original draft, Writing - review & editing.

<u>Dörthe Becker</u>: Conceptualization, Investigation, Resources, Writing - review & editing

<u>Connor S. Murray</u>: Formal analysis, Writing - review & editing

<u>Robert Porter</u>: Investigation, Writing - review & editing

<u>Grace Gutierrez</u>: Investigation, Writing - review & editing

<u>Priscilla Erickson</u>: Investigation, Writing - review & editing

<u>Joaquin C. B. Nunez</u>: Formal analysis, Visualization, Writing - review & editing

<u>Erin Voss</u>: Investigation, Writing - review & editing

<u>Kushal Suryamohan</u>: Formal analysis, Writing - review & editing

<u>Aakrosh Ratan</u>: Formal analysis, Writing - review & editing

<u>Andrew Beckerman</u>: Conceptualization, Resources, Writing - review & editing

<u>Alan O. Bergland</u>: Conceptualization, Data curation, Formal analysis, Funding acquisition, Investigation, Methodology, Project administration, Resources, Software, Supervision, Visualization, Writing - original draft, Writing - review & editing

## Data availability

All scripts and code used for data analysis and plotting are available at https://github.com/alanbergland/DaphniaFromDorset. All sequencing reads have been deposited at NCBI’s Sequence Read Archive (Bioproject # PRJNA725506). The reference genome has been deposited at DDBJ/ENA/GenBank under the accession JAHCQT000000000. Raw phenotype data, the VCF file and associated files used for SNP analysis, as well as Supplemental Table 1 with information on all individuals sequenced are available on Dryad: https://datadryad.org/stash/share/BEppQ_0gXxGRZgsuVmHwwD9T1-0XU4r2YXvpSRlblDc. Maps for Figure 1A are available from (Left panel: https://d-maps.com/carte.php?num_car=2554&lang=en; Middle panel: map taken from OpenStreetMaps; Right panel: Crown copyright and database rights 2021 Ordnance Survey).

## Acknowledgments

The authors acknowledge Research Computing at The University of Virginia for providing computational resources and technical support that have contributed to the results reported within this publication (https://rc.virginia.edu). The authors also thank AnhThu Nguyen for assistance with library construction as well as technical advice and support.

## Supplemental Figures

**Supplemental Figure 1:**
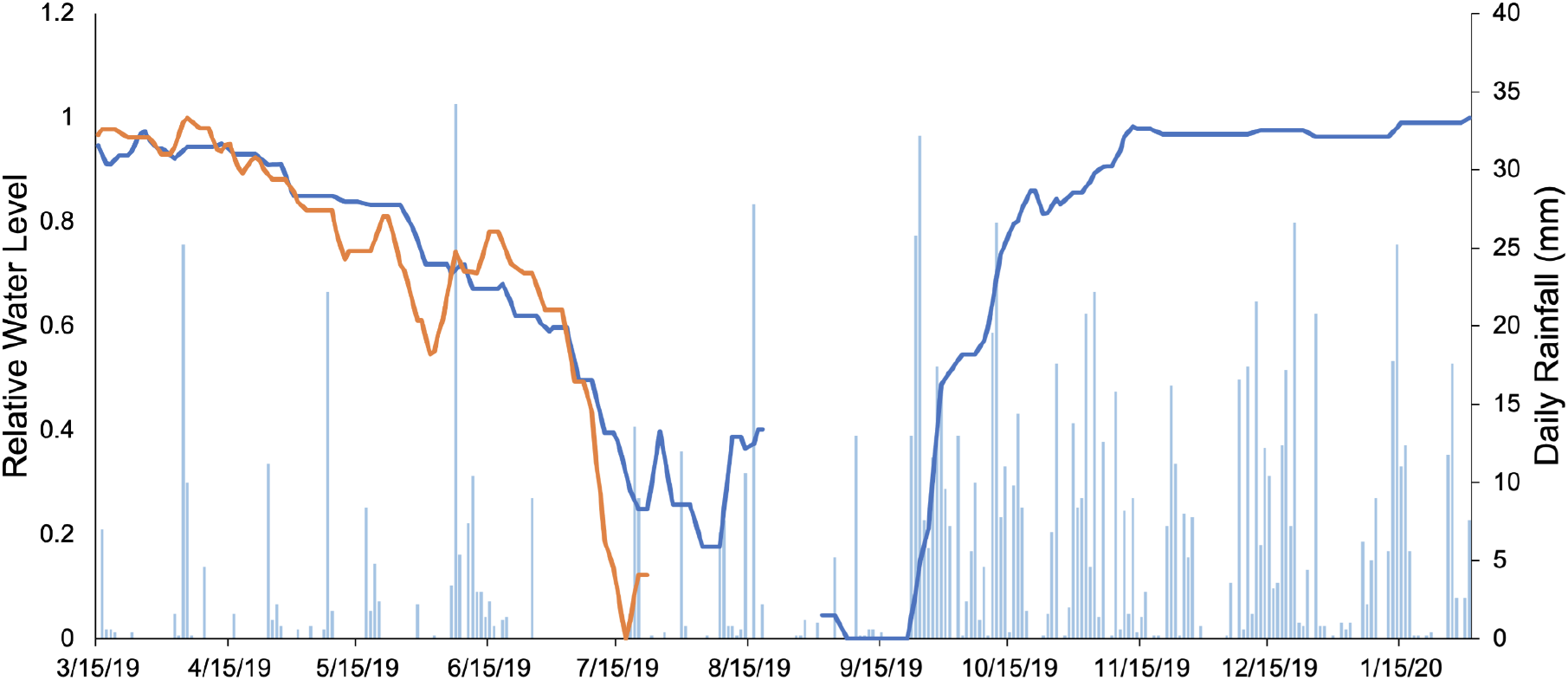
Changes in relative water level in D8 (blue) and DBunk (orange) from March of 2019 through January of 2020 along with rainfall events during the same time period. Ponds are observed to slowly dry during spring and early summer due to decreased rainfall, and then rapidly refill in the fall due to heavy rains. Relative changes in water level were measured from daily photographs taken by time lapse cameras posted at D8 and DBunk. Lowest relative water level measurements do not necessarily correspond to completely dry, but to lowest water level observed.

**Supplemental Figure 2:**
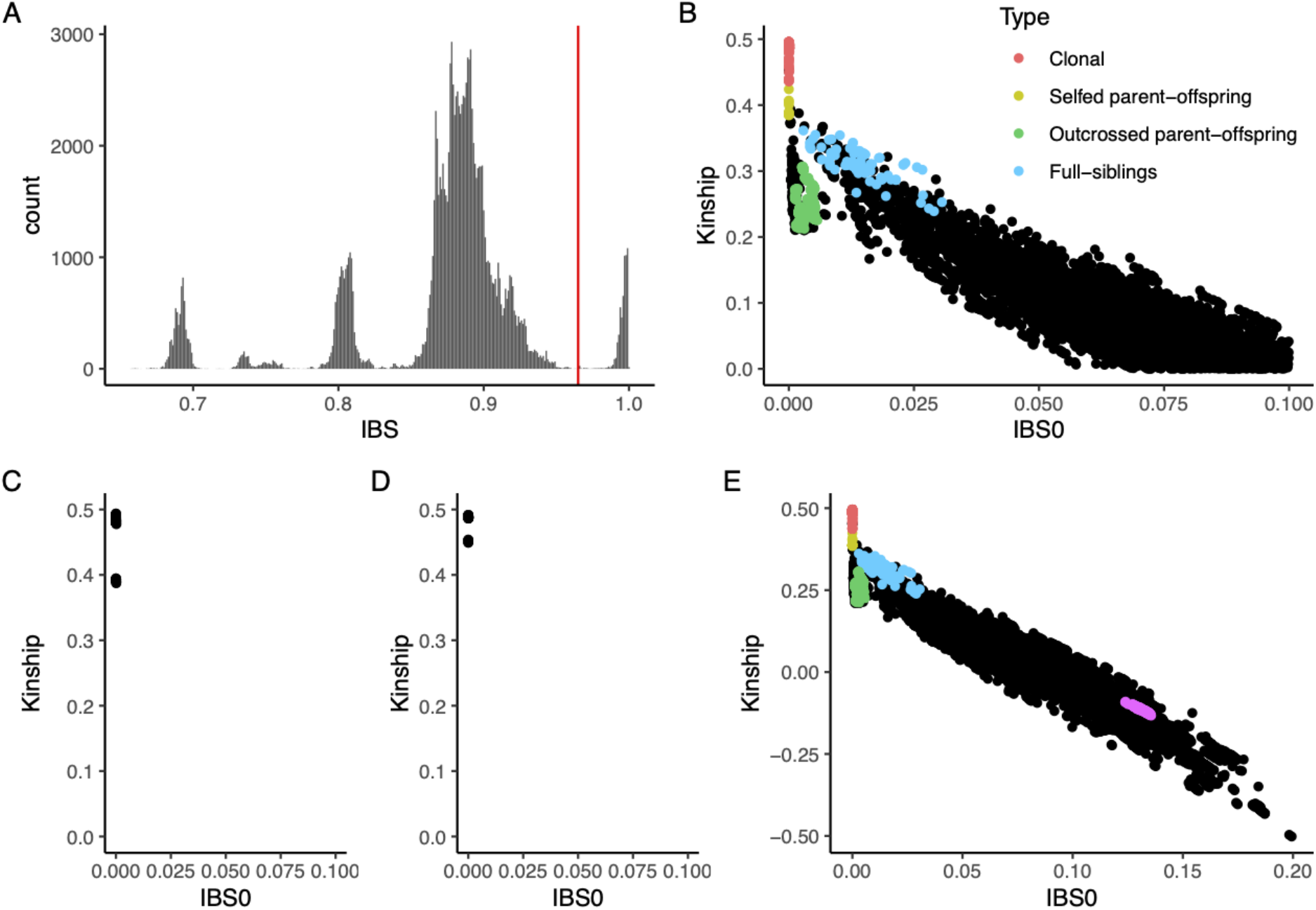
A) Distribution of pairwise IBS values between all sequenced clones with horizontal red line marking the IBS cutoff above which clones were considered to belong to the same clonal lineage. B) Relationship between IBS0 and Kinship as calculated in the program King for pairwise combinations of individuals genotyped from DCat, D8, and DBunk. Only individuals with a median read depth of greater than 14 are included in this graph. Colored points represent prior determined relationships that were used to demarcate clustering and identify additional parent-offspring relationships that were used for constructing the pedigree in Figure 2. C) and D) Examples of outlier individuals within clonal lineages identified by IBS that were subsequently removed from the clonal lineages. E) Zoomed out version of panel B, with A superclone vs C superclone points (in pink) added to illustrate their relationship.

**Supplemental Figure 3.**
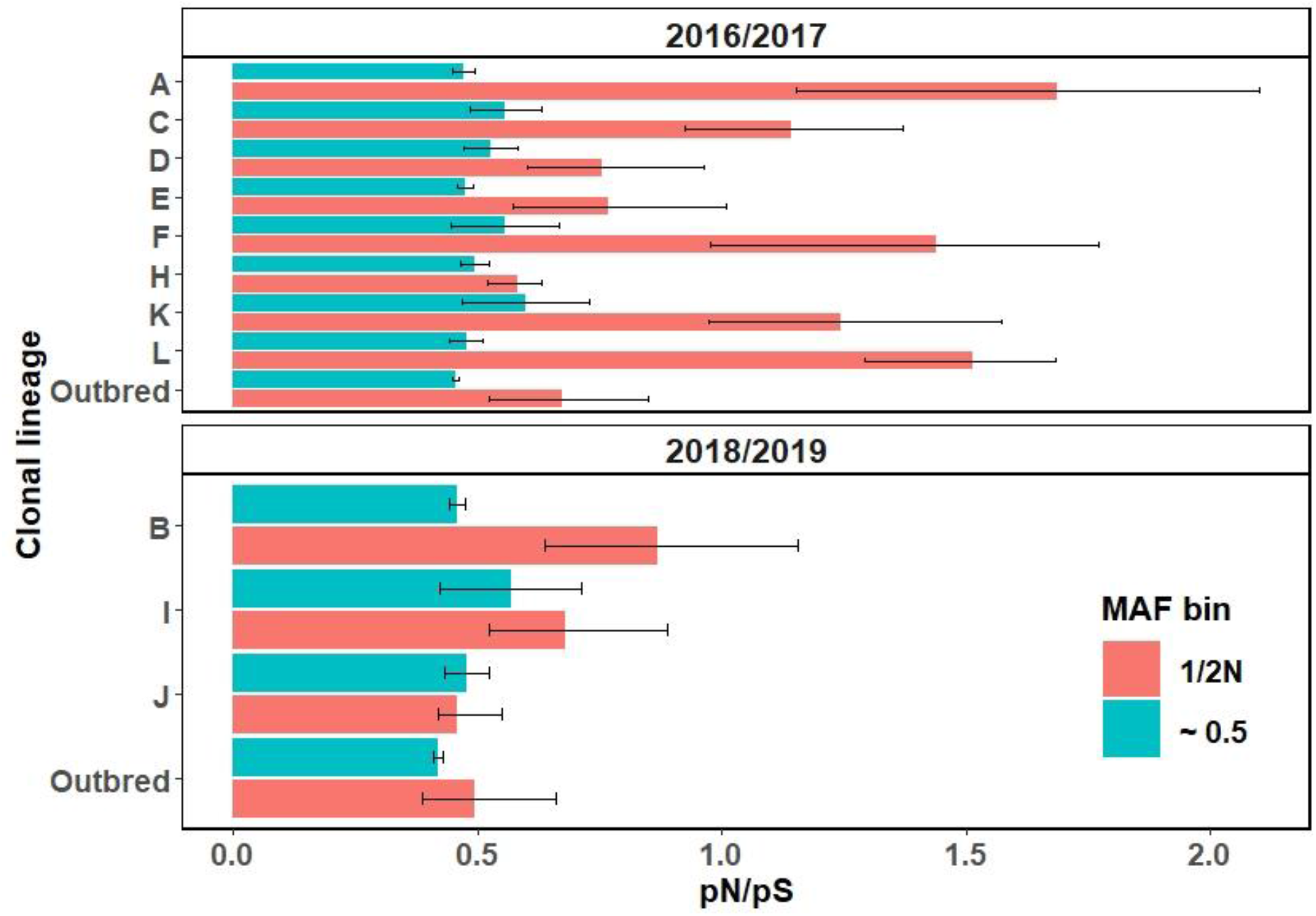
Mutation accumulation. We calculated the ratio of synonymous to non-synonymous mutations at different frequency classes within the larger superclone assemblages (A-L), or among singleton isofemale lines (2016/17) or wild caught individuals (2018/2019). For each group, we randomly selected eight genomes and generated confidence intervals (95%) via bootstrap resampling. See Materials and Methods for details. We identified mutations that were segregating among the sample of genomes, e.g. polymorphism among different field isolates of superclone A. We partitioned those polymorphisms into two classes: New mutations that are segregating at “1/2N” frequency; or, mutations that were shared heterozygotes (“~0.5”). We classified shared heterozygous sites as those where at least 7 (out of 8) individuals were heterozygous. New alleles that arise in asexual lineages are expected to have a higher proportion of non-synonymous sites than older, shared polymorphism. Additionally, the sequencing libraries generated from the 2016/2017 collections were made from isofemale lines that had been propagated in the lab for 4-12 generations, and each library contained ~8-12 individuals. The 2018/2019 libraries were made from single field preserved individuals. The higher pn/ps of the 2016/2017 collections compared to the field caught individuals from 2018/2019 is explainable by these sample features.

**Supplemental Figure 4.**
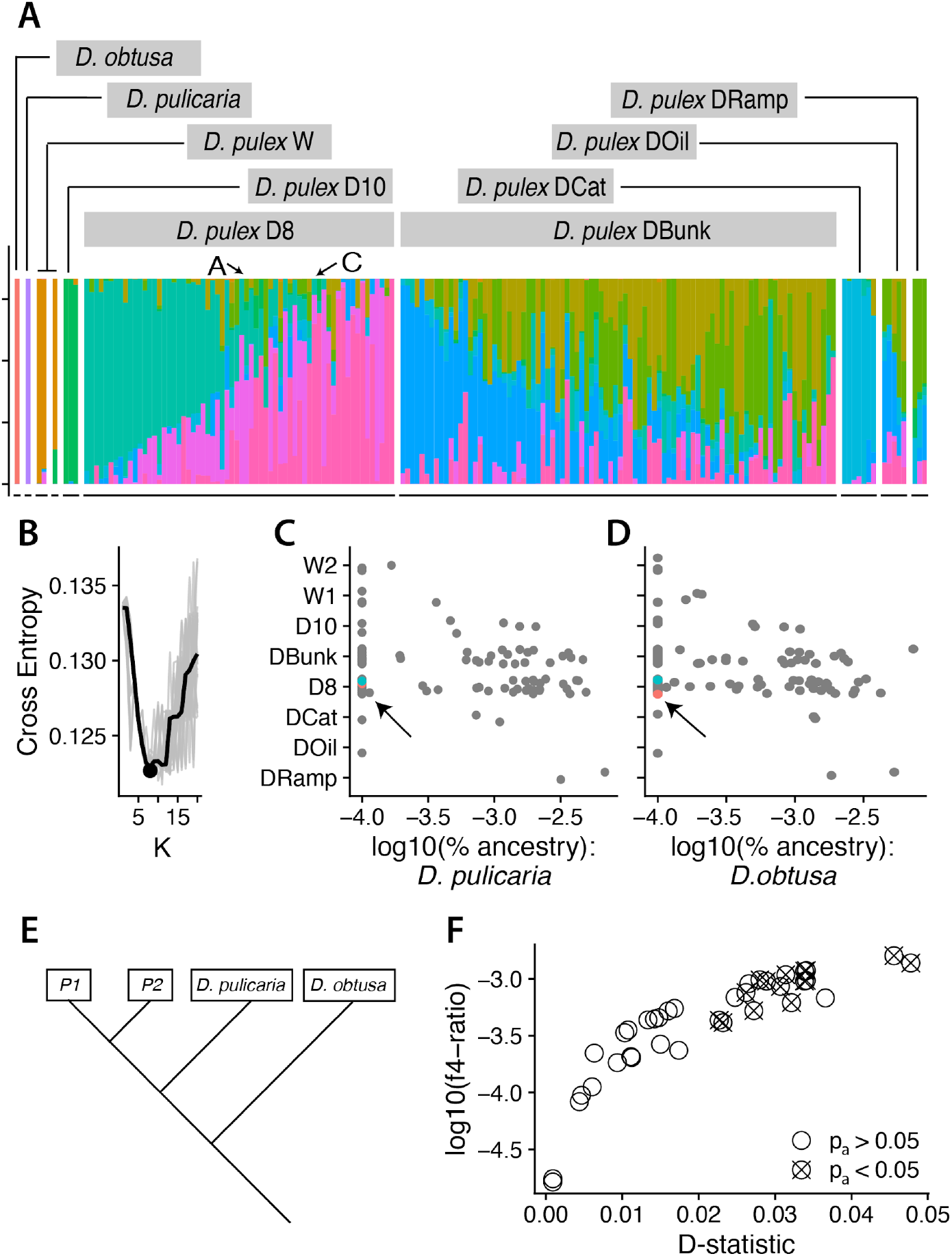
Introgression & hybridization analysis. (A) Structure-type analysis performed using *snmf* function in LEA (see Material and Methods). Each color represents the inferred ancestry coefficient. (B) Cross-entropy analysis suggests that eight clusters is the optimal number of clusters for ancestry analysis. The two out-group species, *D. obtusa* and *D. pulicaria* were assigned to unique clusters, unique clusters were also identified for the Welsh individuals (W) and for individuals sampled from D10. The remaining four clusters were assigned to the Kilwood ponds. (C) and (D) represent the inferred proportion of *D. pulicaria* and *D. obtusa* admixture for the *D. pulex* individuals. Overall, there was a trivial amount of admixture. (E) Schematic of the tree configuration for the *f4*-ratio and D test; P1 and P2 are the population labels for *D. pulex* samples as noted in (A). (F) Several samples show values of D that are statistically different from zero (x-axis), but nonetheless show trivial amounts of ancestry (y-axis).

**Supplemental Figure 5:**
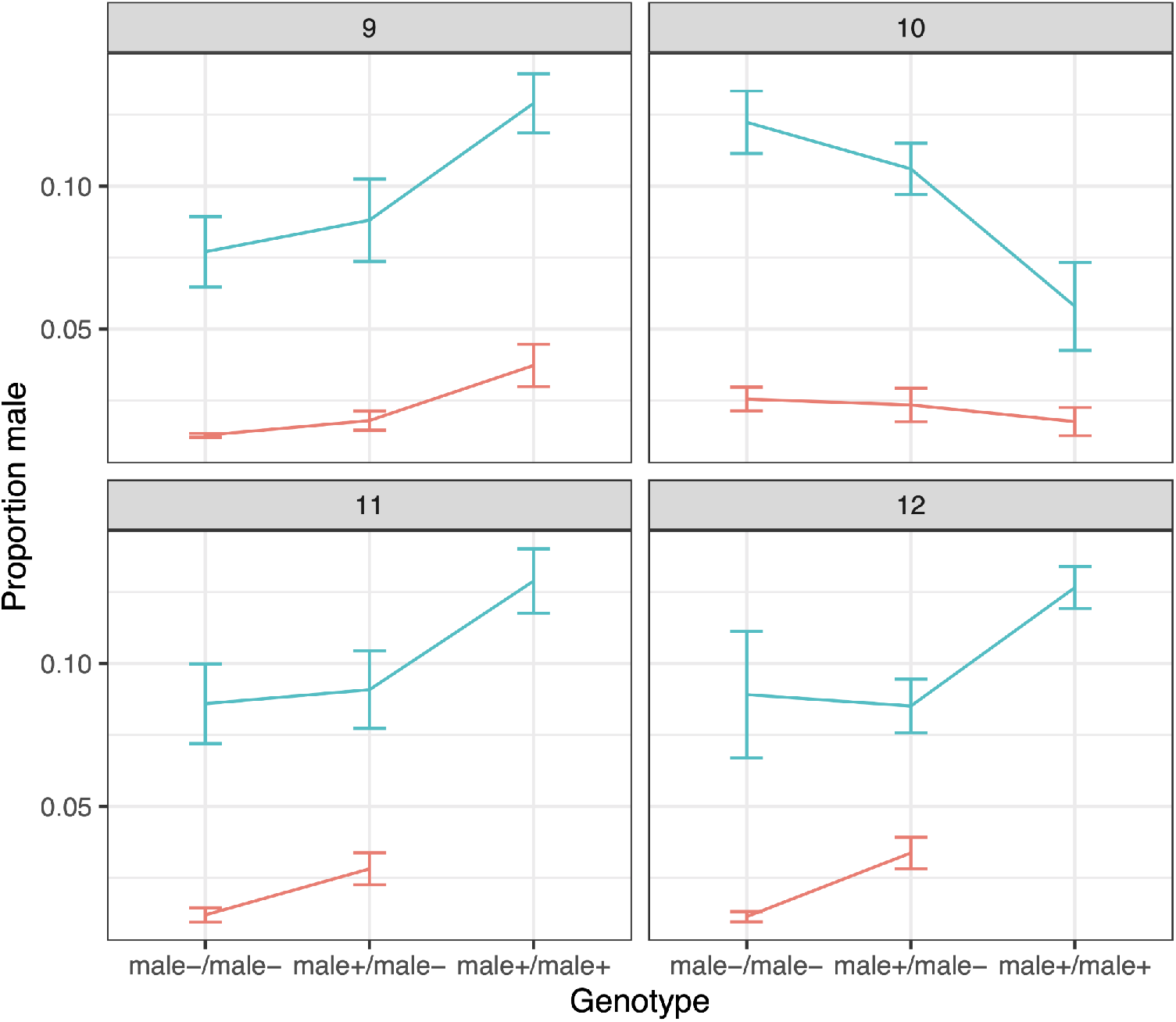
Relationship between genotype and male production for A×C F1 and C×C clones at the 4 pool-seq QTL where genotype had a significant effect on male production (p<0.05). QTL 9 and 12 remained significant after Bonferroni correction. Blue lines represent selfed C clonal lineages, while red lines represent A×C F1 hybrid clonal lineages. Error bars are 95% confidence intervals.

**Supplemental Figure 6:**
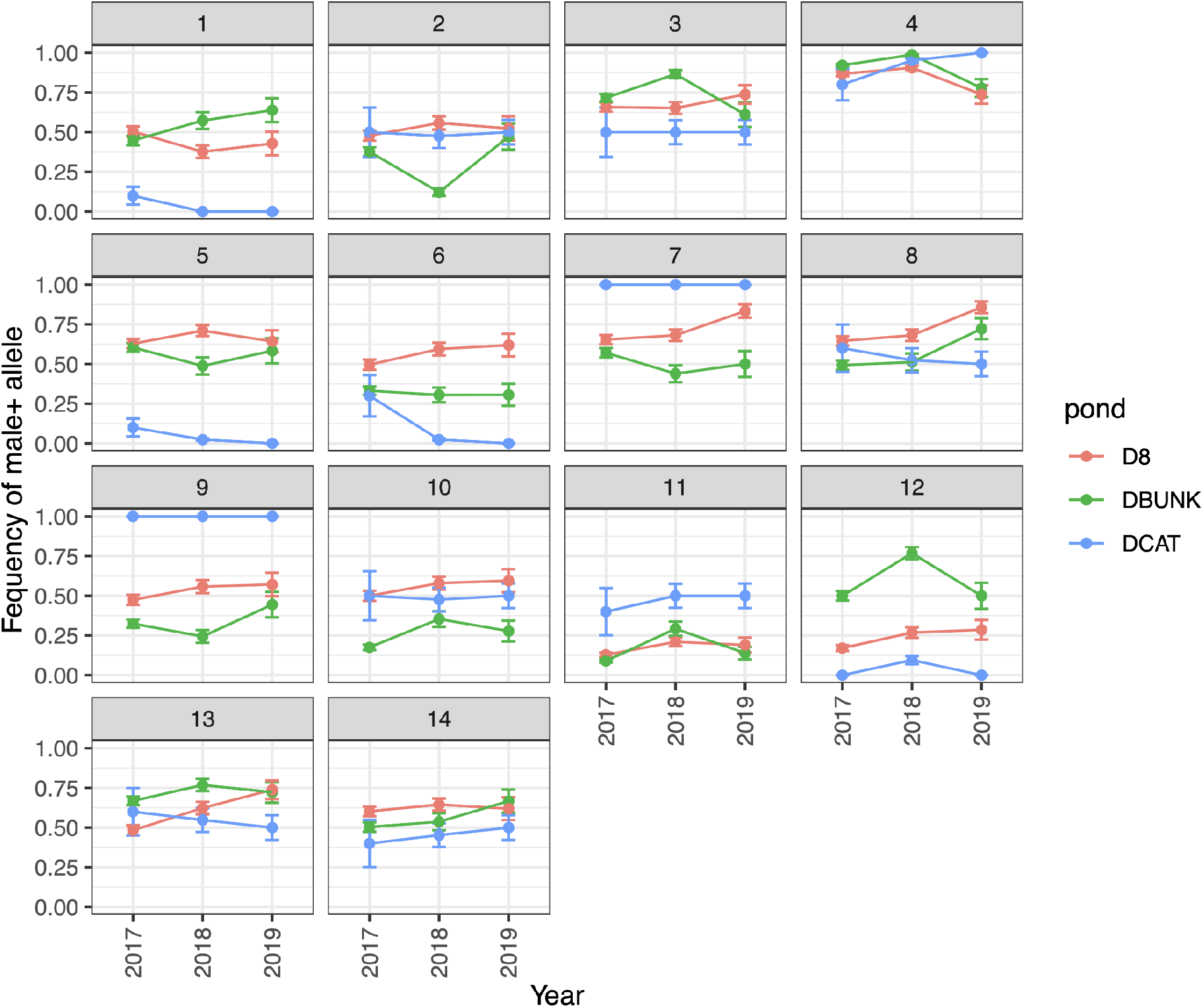
Frequency of *male*^+^ allele at each QTL in the three focal ponds across three sample years. Frequency was calculated across all individuals sampled (no subsetting of superclones). Error bars represent 95% confidence intervals.

**Supplemental Figure 14.**
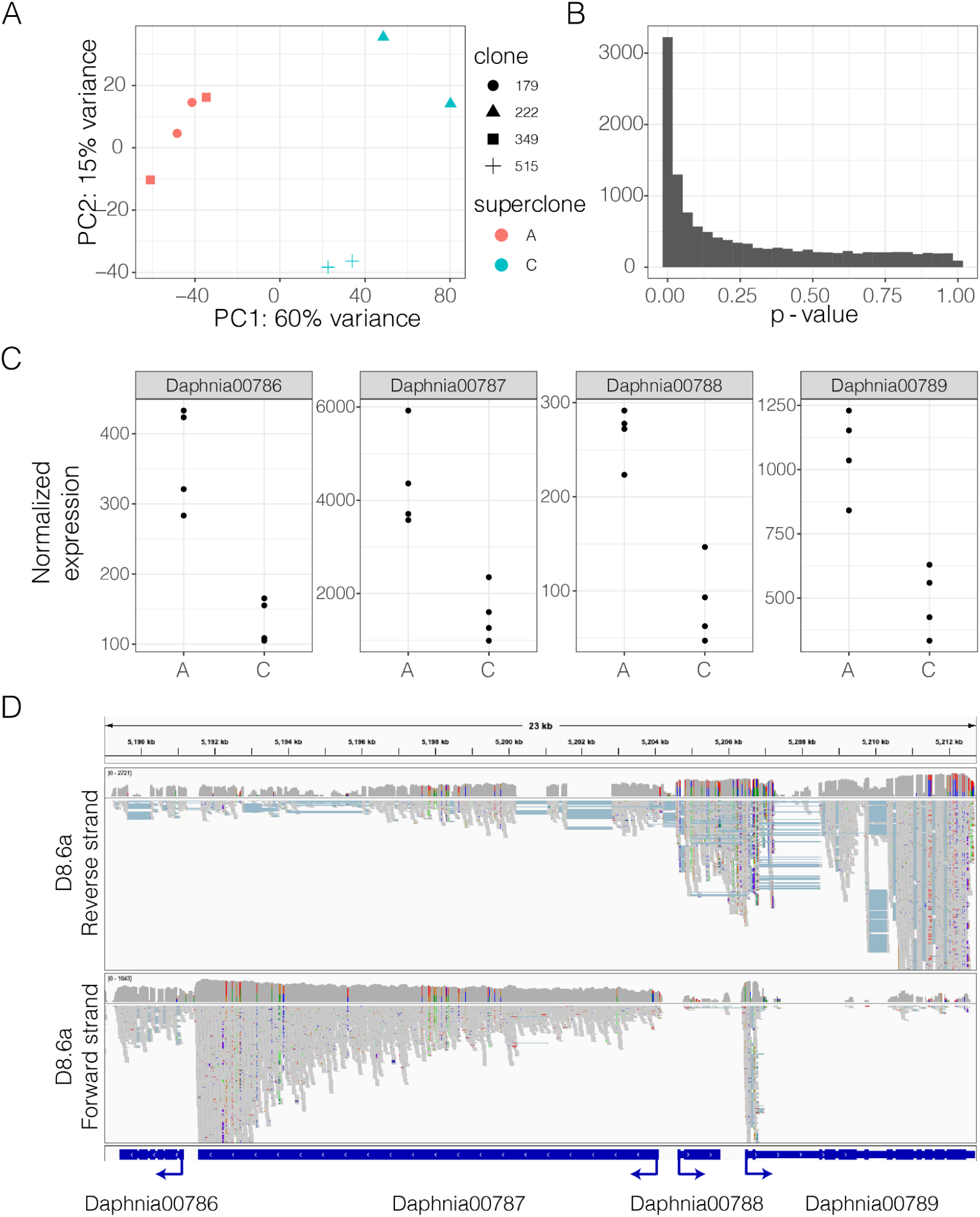
Differential expression between superclones A and C. (A) Principal component analysis shows that PC1 clearly separates superclone A and C, and explains 60% of the variance in gene expression. Clones within a superclone also clustered together. (B) The distribution of p-values contrasting differential expression between superclones. There are many differentially expressed genes. [C] shows the gene expression differences between superclones A and C for the 4 adjacent genes which are strongly differentially expressed. (D) IGView screenshot of the four-gene region from stranded RNA-seq libraries used for gene-model prediction. This screen-shot demonstrates that *Daphnia00787* is intronless, although there are spliced antisense transcripts possibly associated with *Daphnia00788* or *Daphnia00789*.

**Table S1:** List of all genomes sequenced with the following information: NCBI accession number; superclone assignment (OO indicates clonal lineages only sampled one time); sample population; sample year; sample month; median read depth; whether or not the individual was fixed in ethanol in the field (1) or established in the lab prior to sequencing (0, WildSequenced); sex; species; is (1) or is not (0) an AxC F1 hybrid; whether or not the clone was lab generated (the result of crossing in the lab); whether or not the clone was phenotyped in the one liter experiment; and whether or not the clone has low read depth and was excluded from most analyses. https://drive.google.com/file/d/1QRVkgtrxhfMDBtkJNOfD8-Tr6Vwuh-DF/view?usp=sharing

**Table S2:** Clonal diversity for each pond by year. Diversity was calculated using Shannon’s diversity index in the R package *vegan* ^122^.

| Pond | 2016 | 2017 | 2018 | 2019 |
| --- | --- | --- | --- | --- |
| DCat | NA | 1.33 | 0.2 | 0 |
| D8 | 1.39 | 1.02 | 3.05 | 2.91 |
| DBunk | NA | 3.59 | 2.04 | 2.37 |

**Table S3:** List of the 12 D84.A chromosomes, their length, and which North American Daphnia pulex chromosomes they correspond to (PA42, ^69^)

| Chromosome | Length | PA42_Chromosome |
| --- | --- | --- |
| Scaffold_1931_HRSCAF_2197 | 8597654 | 1 |
| Scaffold_9198_HRSCAF_10754 | 12365322 | 2 |
| Scaffold_9199_HRSCAF_10755 | 11497646 | 3 |
| Scaffold_9197_HRSCAF_10753 | 9146106 | 4 |
| Scaffold_9200_HRSCAF_10757 | 10326160 | 5 |
| Scaffold_2373_HRSCAF_2879 | 7726101 | 6 |
| Scaffold_7757_HRSCAF_8726 | 11802501 | 7 |
| Scaffold_6786_HRSCAF_7541 | 13322843 | 8 |
| Scaffold_1863_HRSCAF_2081 | 10446993 | 9 |
| Scaffold_2217_HRSCAF_2652 | 14299301 | 10 |
| Scaffold_9201_HRSCAF_10758 | 6810281 | 11 |
| Scaffold_2158_HRSCAF_2565 | 10306748 | 12 |

